# Prior Experience Alters Exploration in Standard Behavioral Assays More Robustly Than Sex or Light/Dark Cycle in C57BL/6J Mice

**DOI:** 10.64898/2026.11.25.769262

**Authors:** Jake E. Riggi, Margaret H. Copeland, Sydney J. Schaefer, Maya Yun, Jesse D. Cushman

**Affiliations:** Neurobiology Laboratory, National Institute of Environmental Health Sciences, National Institutes of Health, 311 Research Drive, Durham, NC 27710, USA; Neuroscience Program, Lafayette College, 350 Hamilton Street, Easton, PA 18042, USA; Toxicology Program, Department of Biological Sciences, North Carolina State University, 850 Main Campus Drive, Raleigh, NC 27606, USA

**Keywords:** Repeated testing, exploratory behavior, intersession habituation, elevated plus maze, elevated zero maze, open field test, Y-maze spontaneous alternation, novel object recognition

## Abstract

Circadian phase during testing, biological sex, and repeated exposure to the same testing apparatus are all experimental variables known to impact outcomes in neurobehavioral studies. However, the combined, interactive effects of these intrinsic and extrinsic variables on common behavioral tests are not well established, and the effect of repeated testing is particularly understudied. In this study, we examined the compounding impact of these variables by comparing male and female mice on either a standard light cycle or reversed light cycle in the elevated plus maze (EPM), elevated zero maze (EZM), open field test (OFT), Y-maze spontaneous alternation (SA), and novel object recognition (NOR). To examine the impact of repeated exposure to the same testing apparatus, mice were re-exposed to the EPM, EZM, OFT and SA 24 hours after the initial test. We found that exploratory behavior in EPM, EZM, and OFT was robustly diminished on the second exposure. Furthermore, exploration in the open arms of the EPM dropped dramatically, proportionally more than can be explained by the drop in total exploration. The only endpoint unaffected by repeated testing was percent alternation in SA. Light cycle had smaller, more inconsistent effects across assays; male mice on a reversed light-dark cycle showed increased object exploration and improved discrimination in NOR, and mice on a reversed light-dark cycle showed less time exploring the center in the OFT. The effect of sex was minimal, and behavior in EZM and Y-maze was unaffected by sex or light condition. We propose that the first test in any assay elicits an incidental learning experience which is expressed on the second test as a reduction in exploration. The robustness of this test-retest drop in exploration argues that this is a critical experimental variable that must be incorporated into experimental design and interpretation of behavioral results.

## 1. Introduction

Rodent behavioral assays are central tools in neuroscience research as they provide interpretable behavioral measures relevant to cognition, exploration, learning, and threat-avoidance under controlled experimental conditions. These assays have long been used to evaluate pharmacological, neurological, genetic, and environmental impacts on brain function and behavior, and they remain standard approaches for investigating memory, exploration, and threat-related phenotypes (Sousa et al., 2006; Hånell & Marklund, 2014; Crawley, 2008; Lister, 1987; Seibenhener & Wooten, 2015). Performance on behavioral tests is shaped by both intrinsic and extrinsic experimental factors that vary between laboratories. Three such variables include the circadian phase at testing, the sex of the animal, and whether trials are repeated more than once (Hånell & Marklund, 2014; Tsao et al., 2022; Alstott & Timberlake, 2009, Varga et al., 2025). Repeated testing allows for experimental designs that utilize within-subject comparisons, which can reduce the number of animals needed by allowing for multiple manipulations to be tested in the same animal (Festing & Altman 2002; Smith & Lilley 2019). However, the effectiveness of repeat testing depends on whether behavior is sufficiently stable to provide a consistent baseline for comparison.

The potential benefit for reduction of animal numbers means that it is critical to identify what assays can be effectively repeated. Therefore, in this study, we aim to investigate the behavioral changes induced by repeated testing to determine which behavioral tests can be repeated with stable outcomes. We do so by examining five standardized assay paradigms that assess threat avoidance, exploratory behavior, spatial working memory, and object recognition memory: EPM, EZM, OFT, Y-Maze SA, and NOR. We include female and male mice on standard and reversed light schedules and repeat each assay, except NOR, after a 24-hr interval.

The best characterized effect of repeated testing is in the elevated plus maze (EPM), where it is well established that threat-avoidant behavior increases, as indicated by less time spent in the open arms, when rodents have prior experience in the assay (Almeida et al., 1993; Bertoglio and Carobrez, 2000; Schrader et al., 2018; Sergievich et al., 2020; Walf and Frye, 2007; Albernaz-Mariano et al., 2022). This robust effect persists in mice treated with anxiolytics, who also show an increase in threat-avoidance upon repeated testing, despite the continued administration of the anxiolytics (File et., 1993). In the elevated zero maze (EZM), there is conflicting evidence over whether threat-avoidant behavior changes upon repeated testing, though it is generally thought to show more stable performance relative to the EPM (Cook et al., 2002; Tucker & McCabe 2017). In the Y-maze, longitudinal evidence suggests that spontaneous alternation (SA) does not diminish upon repeated trials with large intersession intervals, but the repeatability of this test after a 24-hour delay has not been determined (Cnops et al. 2022; Magen et al., 2012). In the OFT, extensive literature has found that locomotion decreases robustly and consistently upon repeated trials. This phenomenon has been most often studied as intersession habituation, though has also been researched as a useful measure of a learning process (Võikar & Rauvala 2004; Djebari et al., 2026; Nedogreeva et al., 2025; Young et al., 2024; Tzakis et al., 2025; Chen et al., 2023, Wagatsuma et al,. 2018).

A difficulty that arises when studying the effect of repeated testing is that there is no consistent conceptualization of this phenomenon. This lack of consistency is demonstrated by the fact that the effect of repeated testing goes by many names. Depending on the behavioral paradigm, this phenomenon is known as exploratory habituation in the OFT, one-trial tolerance in the EPM, carry-over effects, or novel context recognition (Chen et al., 2023; Heald et al., 2023; Krasne et al., 2015; Wagatsuma et al., 2018).

Another challenge in studying repeated testing is the lack of mechanistic explanation of this phenomenon. There is long-standing evidence that rodents incidentally encode a spatial and contextual representation of novel environments through exploration (Tolman, 1948; Fanselow, 1990; Fanselow, 2000; O’Reilly & Rudy, 2001; Wagatsuma et al., 2018; Leussis & Bolivar, 2006; Krasne et al., 2015; Whittington et al., 2020; Pignatelli et al., 2019). Despite this evidence, the consequences of this incidental learning for routine behavioral phenotyping have rarely been examined as a general process across standard assays. While spatial learning is central to interpretation and explicitly leveraged in some assays like novel object recognition (NOR) and Morris water maze, spatial and contextual learning as general processes may occur in any behavioral test; including those which are typically used to measure anxiety-like behavior, such as the EPM, OFT, and EZM (Chen et al., 2023; Heald et al., 2023; File 1993; Rodgers et al., 1996; Cook et al, 2002; Roy et al., 2009; Wagatsuma et al., 2018; Ferreira et al., 2022; Djebari et al., 2026; Tzakis et al., 2025; Kitamura et al., 2012).

C57BL/6J mice are primarily nocturnal; they are most active during the dark phase and spend much of the light phase sleeping (Ripperger & Albrecht, 2011). Because behavioral testing often occurs during normal working hours, laboratories typically use one of two light/dark schedules. Under a standard 12:12 h light/dark cycle (SLD), lights are on during the day and off at night. This schedule is convenient for researchers, but arousing animals during their rest period disrupts their sleep, which can worsen memory and increase threat-avoidant behavior (Tsao et al., 2022; Schröder et al., 2023; Weber et al., 2024). Under a reversed 12:12 h light/dark cycle (RLD), the light and dark phases are switched, so lights are on at night. During the day, mice are illuminated under dim red light only, which is outside of their visual spectrum (Peirson et al., 2018). This RLD schedule mitigates the need to arouse mice during their rest cycle. The predominant behavioral effect of RLD across assays is that mice move more (Tsao et al., 2022; Richetto et al., 2019). Other behavioral effects of light cycle are assay-specific (Nakano et al., 2016). In this study, we compare behavioral performance between SLD and RLD conditions to determine whether the behavioral changes caused by the circadian phase of testing are influenced by repeated testing and sex.

The last experimental variable under investigation in this study is biological sex, an intrinsic experimental factor which can influence behavioral phenotypes and is important to consider throughout biological research (Clayton & Collins, 2014; Karp et al., 2017; Võikar et al., 2001; Beery & Zucker 2011). Sex can influence endpoints related to most common behavioral assays, as there are sex differences in locomotion, threat-avoidant behavior, learning processes, and circadian regulation (Ryherd et al., 2023; Stevanovic et al., 2022; Bernstein et al., 2024). In addition, sex as a biological variable can be assay and experiment specific, with some studies finding that females spend a greater amount of time in open arm on the EPM and attributing this to females’ greater drive for exploration, while others find that females spend less time in the open arms, citing this as evidence that females have greater threat-avoidant behavior (Baumgartner et al., 2023; Mariscal et al., 2023). In this study, we include both females and males to examine how sex-dependent behavioral patterns vary across SLD and RLD conditions, and repeated testing.

## 2. Methods

### 2.1. Subjects and husbandry

16 female and 16 male 8-week-old C57BL/6J mice (RRID:IMSR, JAX:000664) were obtained from Jackson Laboratory (Bar Harbor, ME, USA). Animals were group housed in standard ventilated mouse cages (Techniplast, Exton, PA) on wood chip bedding (P.J. Murphy Sani-Chip® Bedding, Inotiv, West Lafayette, IN, USA) with additional nesting material (Nestlet-Ancare, Bellmore, NY, USA; Bed-r’Nests-Lab Supply, Missouri City, TX, USA) under constant temperature (70-74° F) and humidity (40-60%) control in an AAALAC-approved animal facility. Mice had *ad libitum* access to standard NIH-31 chow (Ziegler Bros, Inc., Gardners, PA, USA) and reverse osmosis water. Upon arrival, mice were placed into corresponding vivariums to acclimate for 13 days before starting testing. Subjects were identified with individual tail tattoos using a Labstamp automated tail tattoo system (Somark Innovations, San Diego, CA, USA) three days before the start of the first assay. All procedures were approved and performed in compliance with the National Institute of Environmental Health Sciences/National Institutes of Health Humane Care and Use of Animals in Research protocols (Protocol # 2017-0011).

### 2.2. Light phases

Mice were assigned to a standard light/dark cycle (SLD; lights on 7:00-19:00, ZT0=7:00, ZT12=19:00) or a reversed light/dark cycle (RLD; lights off 9:00-21:00 under dim red illumination, lights on 21:00-9:00, ZT0=21:00, ZT12=9:00). RLD mice habituated to reversed light for 11 days before behavioral assays began. Behavioral assays occurred between 9:00-18:00. During behavioral tests, when mice were not in the testing apparatuses, RLD mice were held in a separate dark room with a dim red light lamp, whereas SLD mice were held under normal illumination, at 25 lux.

### 2.3. Mouse handling

Each home cage was provided with a transparent polycarbonate tunnel (2 in wide x 2 in high x 3.5 in long, Mouse Retreat Clear Polycarbonate, BioServ, Flemington, NJ) 5 days before the first behavioral assay to allow mice to acclimate to it. From its introduction onward, mice were exclusively handled with the tunnel to avoid tail handling.

Before each assay, mice were placed in holding rooms adjacent to the testing room for at least 30 minutes. SLD mice were held at 150 lux, while RLD mice were held in the dark and visualized with a red-light headlamp. For each relocation, mice were placed into a transfer cage via a handling tunnel and carried into the testing room, where the assay was performed. When handling mice, experimenters waited until mice freely moved into the tunnel, then closed each side with their hand and lifted the mouse via the tunnel, placing them in a clean transfer cage. This transfer cage was used as a shuttle between the home cage and the behavioral apparatuses, and the handling tunnel was utilized in between each relocation. Each transfer cage corresponded to a single home cage.

### 2.4. Experimental design

These groups were assigned to investigate how circadian timing and biological sex jointly influence behavioral performance. Behavioral testing on animals began after habituation to their light cycle and tunnels. Each behavioral test was followed by a second trial 24-hours later to assess the effect of repeated testing. Animals underwent all trials of the same test within 48 hours, to avoid any effect of day. The sequence of behavioral experiments was EPM, EZM, Y-Maze, OFT, then NOR. Anxiety assays were performed before memory assays, as prior exposure to memory testing could confound subsequent anxiety measures (McIlwain et al., 2001).

### 2.5. Elevated plus maze and elevated zero maze

The EPM (catalog no. 60140; Stoelting Co., Wood Dale, IL, USA) consists of a raised maze 60cm in height in the shape of a plus, with two open arms, two closed arms and a center point in the middle of the maze. The EZM (Noldus, Wageningen, the Netherlands) consists of a raised circle 61cm in height separated into four quadrants, two open quadrants and two closed quadrants perpendicular from each other. Mice were placed into the center portion of the EPM, or towards a closed quadrant of the EZM. The room was indirectly lit with the open arms/quadrants measuring 150 lux. EZM had custom 3D printed walls 5mm high on the open quadrants to prevent mice from falling. However, 6 mice were tested before the custom walls were printed, and 2 fell off of the apparatus onto a black mat cleaned with Pre-empt. Thus, these 6 mice were replaced in the EZM analysis. For both EPM and EZM, the center-point of the mice were tracked via EthoVision XT 19. The total distance moved, time spent in open and closed arms (as well as center for the EPM), and the distance moved within the open and closed arms, was measured. Recording began 7 seconds after a center-point was detected, to allow experimenters time to leave the field of view of the camera behind a black curtain. If experimenters remained in the field of view after 7 seconds, the data in which the experimenter remained visible was excluded from analysis. The apparatus and mat were cleaned with Pre-empt and dried before and after every day of trials, and cleaned with isopropyl alcohol between each trial.

### 2.6. Spontaneous alternation in Y-maze

The standard Y-maze (Med Associates Inc.) consisted of three identical arms (15 in W × 15 in L × 7.8 in H) facing towards each other in the shape of the letter “Y”. Distal cues in the shape of a star, pentagon, plus sign, and triangle were placed on each wall of the testing room held at 35 lux to assist with the subjects’ orientation. Mice were consistently placed in a specific arm of the maze and were allowed to explore the maze freely. The Y-maze was cleaned with 70% isopropyl alcohol between trials and cleaned with Pre-empt at the beginning and end of all testing days.

The center points of the mice were tracked using Ethovision XT19. A spontaneous alternation was scored as a mouse entering the arm in which they had not previously explored, entering three distinct arms sequentially. The “percent alternation” was calculated by taking the amount of correct alternations divided by the total number of alternations.

### 2.7. Open field test and novel object recognition

The combined OFT and NOR were conducted over four consecutive days, with one 10-minute trial per mouse each day, using four opaque white Noldus PhenoTyper home-cage observation boxes (3000 series; Noldus, Wageningen, the Netherlands; 30 × 30 × 35 cm [L × W × H]). One mouse was tested per box under 35 lux illumination. Behavior was recorded via a ceiling-mounted infrared camera within the PhenoTyper observation boxes. Mice were brought into the testing room one cage at a time and tunnel handled into and out of the boxes. Each mouse was placed in the center of the observation box; placement order depended on which mouse entered the handling tunnel first. After video recording began, tracking started automatically 5 seconds after each mouse was placed in a box, allowing the handler time to remove the handling tunnel. After each trial, mice were returned to their home cage in placement order. Home cages were returned directly to the vivarium after all mice had been retrieved. The Phenotyper boxes and objects were cleaned with Pre-Empt before the next cage was brought into the room.

Days 1 and 2 constituted the two-day OFT and simultaneously served as habituation to the chambers for NOR. The tracking arena was calibrated using its 30 cm side length. For OFT analysis, the center zone area was defined as 50% of the arena total, forming a 21.21 x 21.21 cm square; the remaining area was designated as the periphery. The total distance traveled, the percentage of time spent in the center zone, the number of entries into the center zone and the latency to the first entry to the center zone were examined.

NOR was conducted on Days 3 and 4. Two object types, either a 25 mL Erlenmeyer flask containing a wad of paper with the opening secured by a black marble, or a 20 mL vial containing water, were used (Supp 8). These object types were chosen as they were similar in size and material but different in shape and appearance. The type of object used during familiarization was counterbalanced across the four boxes to account for possible object preferences. On Day 3 (familiarization), two identical objects of either type, labeled as “familiar” objects, were placed in the upper left and right corners of each chamber and secured with strong adhesive double-sided tape down to prevent getting knocked over. On Day 4 (test), one familiar object in each box was replaced with the other object type, deemed as the “novel” object. Circular investigation zones around the objects were sized to detect nose-point entry near each object while excluding unrelated nose movements. Nose-point investigation for object preference was measured by automated EthoVision analysis, which was validated against manual scoring (Supp 10). A gap between the walls of one PhenoTyper box across all four days confounded our data by amplifying exploration in this region. Therefore, all data collected from mice tested in Box 3 were excluded from analysis.

### 2.8. Data analysis and statistics

Ethovision XT 19 was used to track mice, identify their position relative to manually set zones of the apparatuses, and perform preliminary data analysis and scoring of pertinent behavioral endpoints. Across all assays, behavior was recorded via an infrared camera. For the Y-maze, OFT, and NOR, scoring and tracking were manually validated using behavioral videos for 1-3 hours per assay to ensure accuracy of automated methods. With exported data from Ethovision, JMP (Version 18.2.2) and R Studio (Version 4.4.1) were used for further data analysis and statistics. R Studio was used for data visualization.

Statistical tests for EPM, EZM, Y-maze, OFT, and for total object investigation in NOR began with a two-factor repeated measures ANOVA, with sex and light condition as between-subject variables and day as a within-subject variable. If a significant or trending (p<0.1) effect of sex, light condition, day, or an interaction thereof was identified, a Bonferroni test was performed to compare the groups of interest (Supp 11). Moreover, when an ANOVA showed a significant effect, mice of the group of interest were pooled across other variables, and a student’s t-test was performed to characterize the difference between these groups (Supp 12-13). For NOR, statistical tests for percent time investigating novel object and discrimination ratio began with a two-factor ANOVA, with sex and light condition as between-subject variables. Percent novel investigation was compared to percent familiar investigation using a repeated measure of object (Supp 14). Discrimination ratio was assessed using a two-sided one-sample t-test to identify discrimination ratios that were significantly different than zero (Supp 15).

## 3. Results

### 3.1. EPM exploration is diminished by repeated testing, and closed arm exploration is reduced in males

In EPM, distance moved during the trial was measured to assess locomotor activity as a measure of exploratory behavior (Fig 1, Supp 1, Supp 2). Four groups were compared: female SLD (n=8), male SLD (n=8), female RLD (n=8), and male RLD (n=8). There was a trend towards females moving more in the maze than males (p_sex_=0.0761, Fig 1C). Then, the percentage of time of the trial that mice spend in the open arm, closed arm, and center was quantified to assess threat-avoidant behavior (Fig 1B, Supp 1). Percent time in open arms remained stable across sex and light condition (p_sex_=0.7630, p_light_=0.8218). There were no interactions between day, sex, or light condition in distance moved or percent time in open arms.

**Figure 1:**
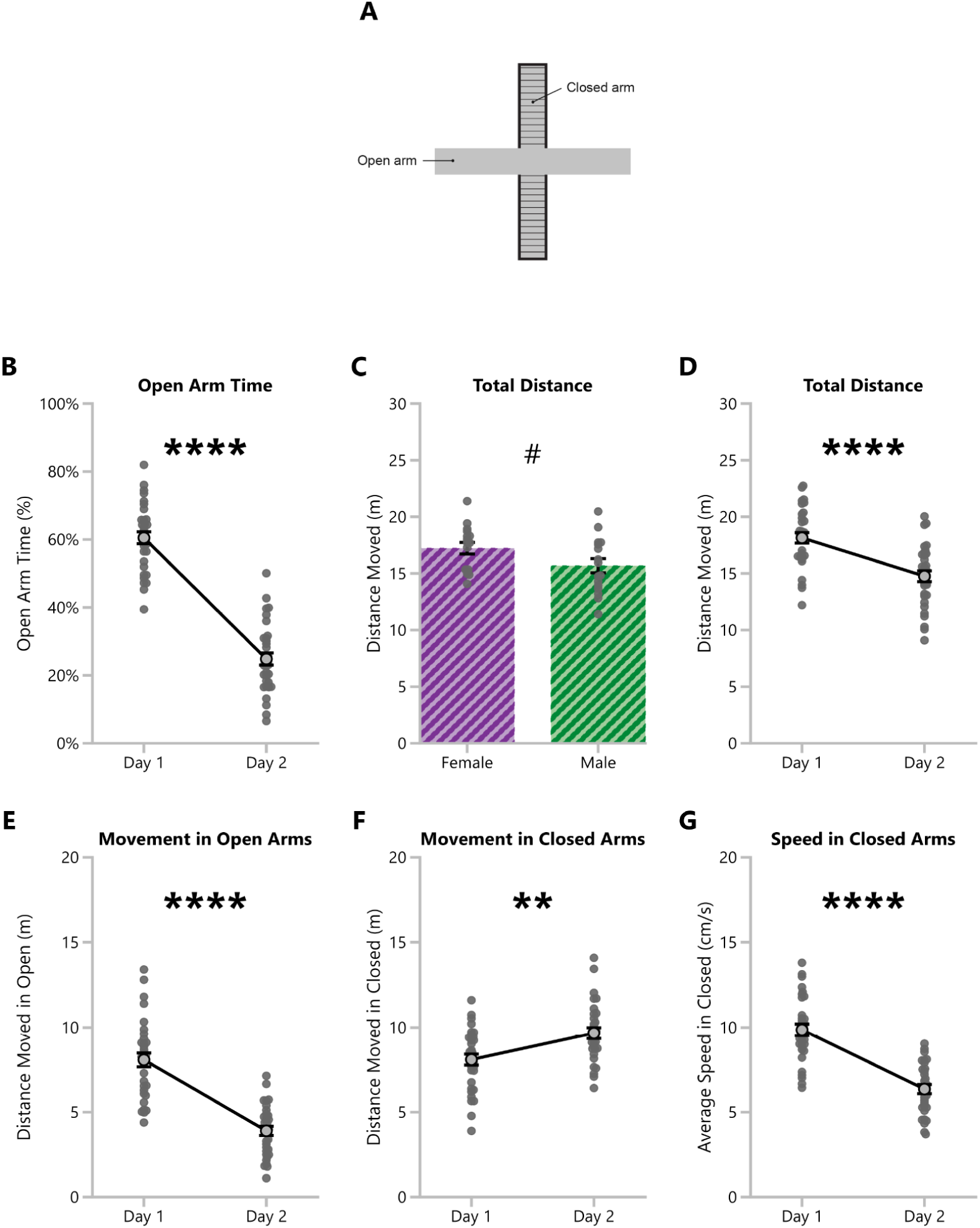
EPM closed arm exploration is reduced in males, and EPM open arm exploration is diminished by repeated testing. **A:** Diagram of elevated plus maze. **B:** The percent of time in open arms for all mice on day 1 compared to repeated testing on day 2. Mean and standard error are shown. **C:** The total distance moved over the duration of a trial in meters for female and male mice. Data points represent the average of both days for each mouse. Mean and standard error are shown. **D:** The total distance moved over the duration of a trial for all mice on day 1 compared to repeated testing on day 2. Mean and standard error are shown. **E:** The total distance moved in open arms only over the duration of a trial for all mice on day 1 compared to repeated testing on day 2. Mean and standard error are shown. **F:** The total distance moved in closed arms only over the duration of a trial for all mice on day 1 compared to repeated testing on day 2. Mean and standard error are shown. **G:** The average speed in closed arms in centimeters per second for all mice on day 1 compared to repeated testing on day 2. Mean and standard error are shown. ^#^p<0.1, *p<0.05, **p<0.01, ***p<0.001, ****p<0.0001.

The greatest effect in EPM were behavioral changes during the second day compared to the first. On the second test, percent duration in the open arms was reduced by 24% (p<0.0001, Fig 1B) compared to the initial assay. Repeated testing decreased open arm percent for all groups, regardless of sex or light condition (p_sex_=0.8388, p_light_=0.8254, Supp 1). Repeated testing also decreased locomotor activity; on day 2, subjects moved 16% less (p<0.0001, Fig 1D). These reductions in open arm time and locomotion indicate that exploration decreases and threat avoidance increases upon re-exposure to EPM, regardless of light condition and sex.

Then, we examined the distance moved within the open and closed arms individually. Distance moved within the open arms dropped significantly after repeated testing (p<0.0001, Fig 1E), whereas distance moved in closed arms increased significantly on day 2 (p=0.0008, Fig 1F). These data indicate that the overall drop in the total distance moved after repeated testing is driven by a decrease in locomotion within the open arms, despite an increase in closed arms locomotion. Moreover, we observed a trend towards a sex difference in locomotion in the closed arms (p_sex_=0.0725), but not in the open arms (p_sex_=0.4294). Investigating this sex difference trend in closed arm behavior, we also observed that females move faster in the closed arms than males (p_sex_=0.0333). There is a significant day*sex effect (p_day*sex_=0.0472, Fig 1G) in speed in the closed arms.

### 3.2. EZM exploration is diminished by repeated testing

Behavior was also examined in the EZM, including distance moved and percent time in open (Fig 2, Supp 3, Supp 4). Compared to EPM, RLD light condition showed a greater effect on locomotion. Distance moved was specifically increased for RLD male mice compared to SLD male mice (p=0.0473, sex*light interaction; p_light_=0.0644 for males only, p_light_=0.3020 for females only, Fig 2B). There were no sex differences in locomotion. Moreover, similar to EPM, percent time in open arms remained stable across light conditions and sex (p_light_=0.2323, p_sex_=0.6768). Mice in EZM did show reduced open arm time (p=0.0185, Fig 2C) and distance moved (p=0.0101, Fig 2D) on day 2 compared to day 1, but this second-trial reduction was less dramatic than in EPM.

**Figure 2:**
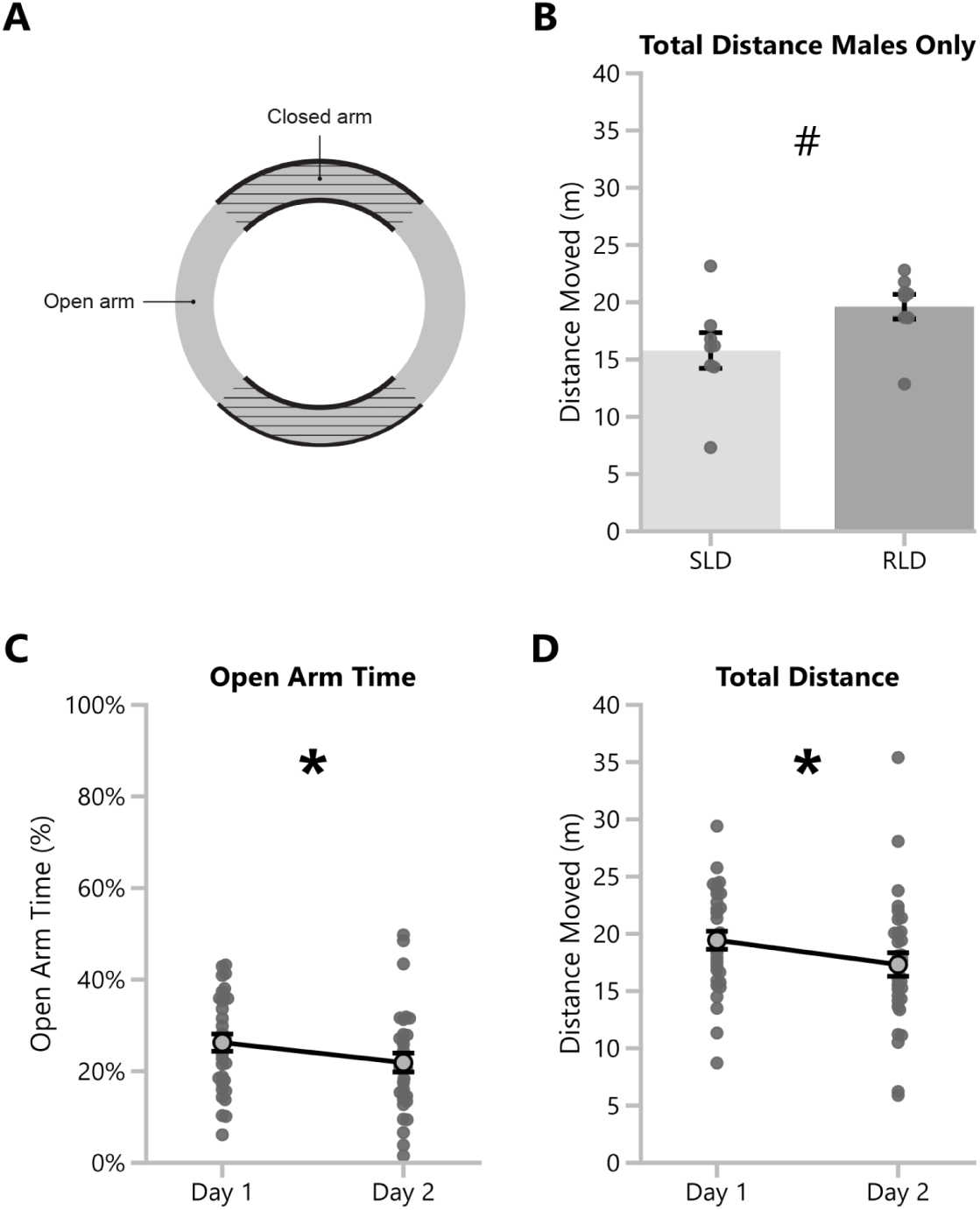
EZM exploration is diminished by repeated testing. **A:** Diagram of elevated zero maze. **B:** The total distance moved over the duration of a trial for male reversed light-dark cycle and male standard light-dark cycle mice. Data points represent the average of both days for each mouse. Mean and standard error are shown. **C:** The percent of time in open arms for all mice on day 1 compared to repeated testing on day 2. Mean and standard error are shown. **D:** The total distance moved over the duration of a trial for all mice on day 1 compared to repeated testing on day 2. Mean and standard error are shown. ^#^p<0.1, *p<0.05, **p<0.01.

### 3.3. Y-maze exploration, but not spontaneous alternation, is reduced by repeated testing

The primary endpoint of the Y-maze is spontaneous alternation (SA), a measure of spatial working memory (Fig 3, Supp 5). A correct SA is completed when a mouse, when faced with the choice of entering a new arm, enters the one less recently visited. Mice with impaired spatial working memory struggle to determine which arm they have most recently visited, thus show deficits in SA (Kraeuter et al., 2019; Hughes, 2004). We observed no effect of sex or light condition on SA (p_light_=0.9982, p_sex_=0.2983), distance moved (p_light_=0.4283, p_sex_=0.9099), or total arm visits (p_light_=0.5594, p_sex_=0.5211). When the assay was repeated 24 hours later, distance moved was significantly reduced on day 2 (p<.0001, Fig 3D), with a trend towards a day*light cycle interaction on distance moved (p_day*light_=0.0731). Similarly, total visits were significantly reduced on day 2 (p<0.001, Fig 3C), with a trend towards a day*light cycle interaction (p_day*light_=0.0851). These data indicate repeated testing influences behavioral performance in the Y-Maze. However, SA was unchanged on the second day (p_day_=0.1304, Fig 3B), indicating reduced exploratory behavior but maintained spatial working memory on the second trial. These data indicate if the intended readout of Y-maze is SA, this is a repeatable test, unlike EPM, EZM, or OFT.

**Figure 3:**
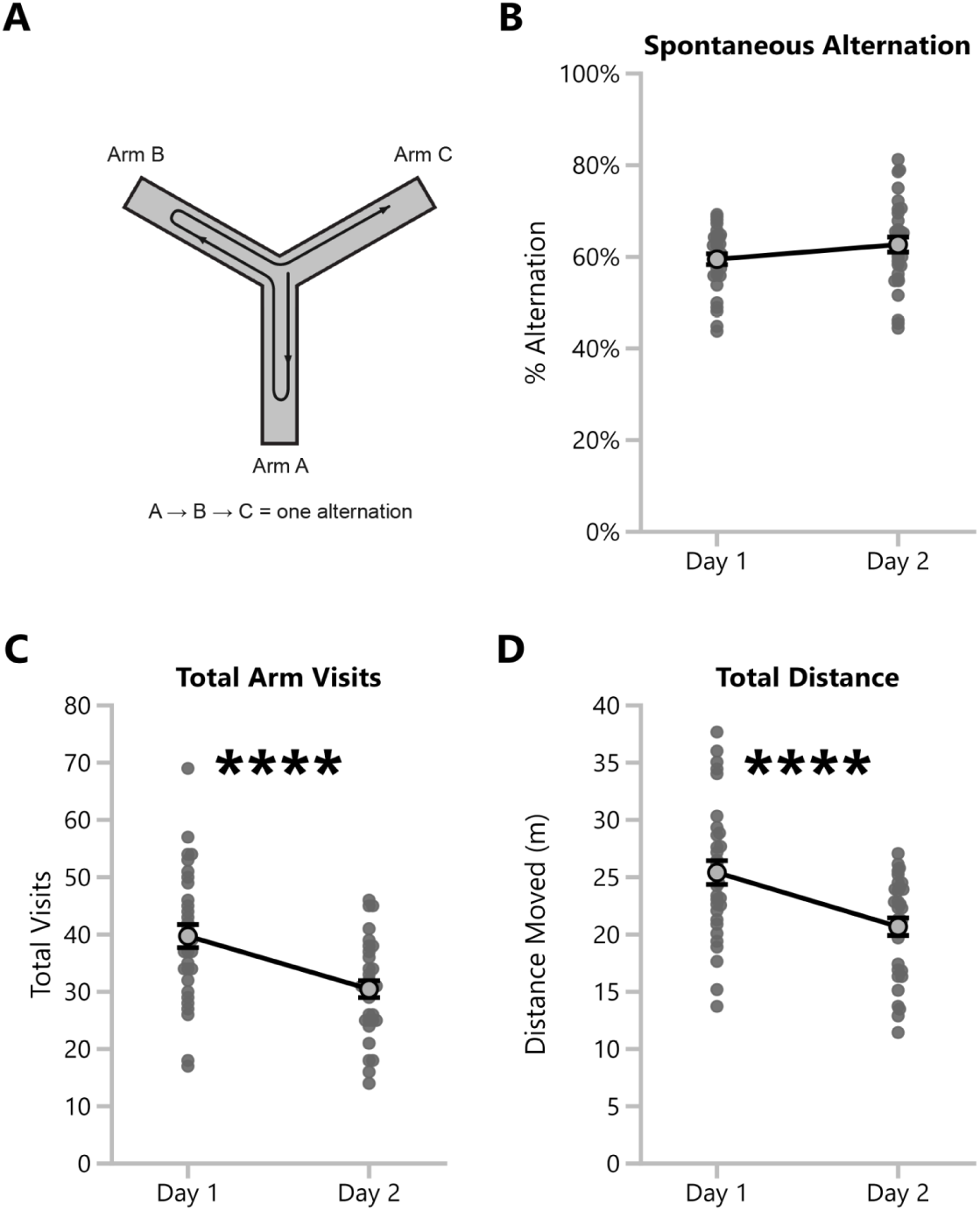
Y-maze exploration, but not spontaneous alternation, is diminished by repeated testing. **A:** Diagram of Y-maze. **B:** The percent alternation for all mice on day 1 compared to repeated testing on day 2. Mean and standard error are shown. **C:** The total arm visits for all mice on day 1 compared to repeated testing on day 2. Mean and standard error are shown. **D:** The total distance moved over the duration of a trial for all mice on day 1 compared to repeated testing on day 2. Mean and standard error are shown. ****p<0.0001.

### 3.4. OFT center time is reduced by reversed light-dark cycle, and OFT exploration is diminished by repeated testing

Mice were tested on the OFT to quantify their time in the center, which is considered a more threatening area, compared to the periphery (Fig 4, Supp 6, Supp 7). The percent of time during the test in the center was significantly lower for mice on the reverse light-dark schedule (p_light_=0.0447, Fig 4A). However, these results do not indicate that RLD mice have reduced exploratory behavior as there was a trend towards RLD having overall higher locomotor activity compared to SLD mice (p_light_=0.0866, Fig 4C).

**Figure 4:**
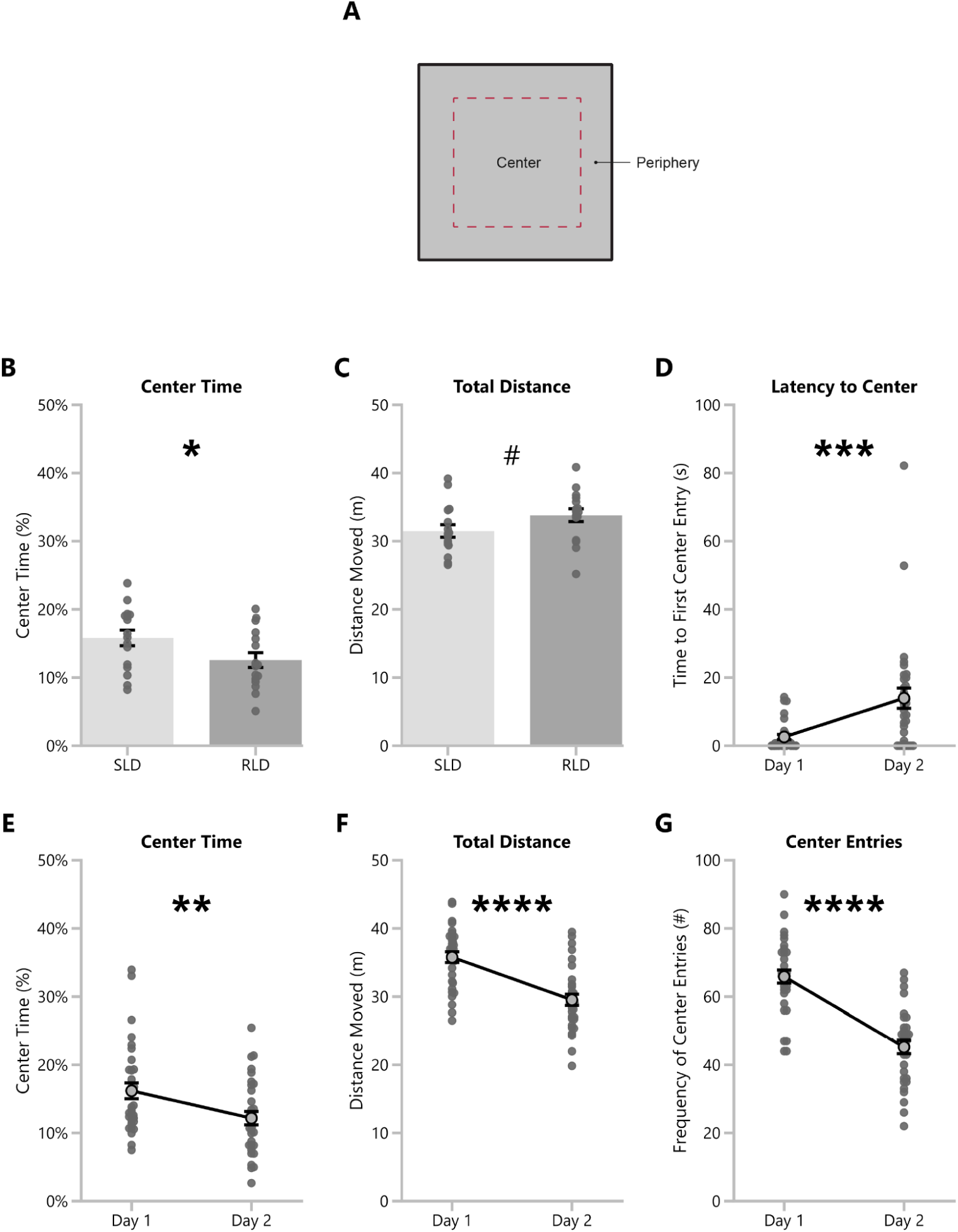
OFT center time is reduced by reversed light-dark cycle, and OFT exploration is diminished by repeated testing. **A:** Diagram of open field test. **B:** The percent of trial time in the center of the open field for reversed light-dark cycle and standard light-dark cycle mice. Data points represent the average of both days for each mouse. Mean and standard error are shown. **C:** The total distance moved over the duration of a trial for reversed light-dark cycle and standard light-dark cycle mice. Data points represent the average of both days for each mouse. Mean and standard error are shown. **D:** The time in seconds from the start of the trial to the first entry to the center for all mice on day 1 compared to repeated testing on day 2. Mean and standard error are shown. **E:** The percent of trial time in the center of the open field for all mice on day 1 compared to repeated testing on day 2. Mean and standard error are shown. **F:** The total distance moved over the duration of a trial for all mice on day 1 compared to repeated testing on day 2. Mean and standard error are shown. **G:** The total number of entries to the center of the field from the periphery over the duration of a trial for all mice on day 1 compared to repeated testing on day 2. Mean and standard error are shown. ^#^p<0.1, *p<0.05, **p<0.01, ***p<0.001, ****p<0.0001.

On the second trial, the percent of time in the center (p_day_=0.0075, Fig 4B) and distance moved (p_day_<0.0001, Fig 4B) were both reduced, indicating reduced exploration after repeated testing. Consistent with diminished exploratory behavior on day 2, latency to center was increased on day 2 (p_day_= 0.0006, Fig 4E), and the number of center entries was reduced (p_day_<0.0001, Fig 4F). Center time, distance moved, and latency to center were unaffected by sex, and there were no statistical interactions between light, sex, or day in any of these measures.

### 3.5. Object exploration is increased by reversed-light cycle, and NOR performance is best in reversed-light cycle males

Finally, cognition and memory were assessed in NOR. Overall, RLD mice spent more time investigating the objects (p=0.0353, Fig 5A), indicating an overall increase in exploratory behavior. There was a trend towards a day*sex*light interaction (p= 0.0835) for total object investigation, and a Bonferroni analysis revealed that on the familiarization day, males exhibited a difference between male SLD and RLD conditions, where male RLD mice explored the objects significantly more than male SLD mice (p=0.001, Fig 5B). This increased exploration for male RLD mice compared to male SLD mice on day 3 is correlated with increased recognition of the novel versus familiar object on day 4 in male RLD mice (p=0.014, Fig 5D). However, there was no significant or trending effect of sex or light on percent novel investigation or discrimination ratio between familiar and novel objects.

**Figure 5:**
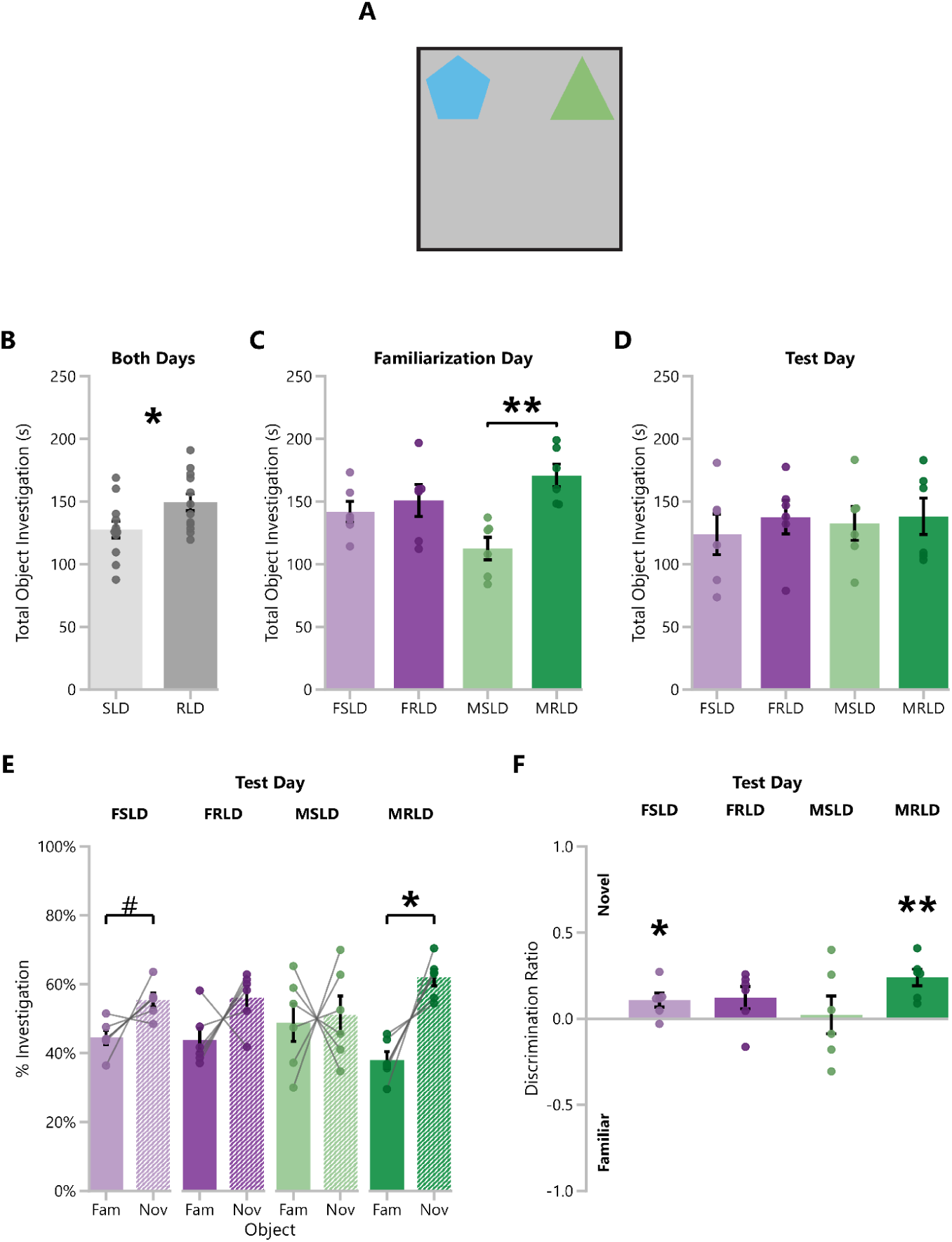
Object exploration is increased by reversed-light cycle, and NOR is improved in male mice on reversed-light cycle compared to standard light-cycle mice. **A:** Diagram of novel object recognition test. **B:** The total time in seconds of object investigation for reversed light-dark cycle and standard light-dark cycle mice. Data points represent the average of both days for each mouse and includes investigation of both novel and familiar objects. Mean and standard error are shown. **C-D:** The total time in seconds of object investigation for each of the four experimental groups on the familiarization day (**C**) and test (**D**) day. Mean and standard error are shown. **E:** Percent of object investigation time allotted to the novel object and the familiar object for each of the four experimental groups on the test day. Mean and standard error are shown. **F:** Discrimination ratio between the novel and familiar object for each of the four experimental groups on the test day. Mean and standard error are shown. DR=1 indicates complete preference towards the novel object, DR=-1 indicates complete preference towards the familiar object, and DR=0 indicates equal preference towards each object. FLD = Female standard light/dark, MLD = male standard light/dark, FRLD = Female reversed light/dark, MRLD = male reversed light/dark, ^#^p<0.1, *p<0.05, **p<0.01, ***p<0.001, ****p<0.0001.

To investigate whether mice identified the novel object over the familiar object, we first examined the percent of object investigation which was allotted towards the novel object compared to the percent allotted towards the familiar object (Fig 5E). We adjusted exploration of the novel object for total investigation so that the results are not confounded by differences in the total exploration towards the objects. Comparing the novel and familiar investigation, we found that male RLD mice showed the largest difference between novel and familiar exploration (p=0.014, Fig 5E). Due to a limited sample size, female RLD and SLD mice did not significantly show a higher percent novel investigation compared to familiar when examined independently. However, when the female RLD and SLD mice were pooled together, they also showed a significant recognition of the novel object over the familiar one (p=0.0257). Then, we examined the recognition of the novel object using a discrimination ratio (Fig 5F). Similarly, the male RLD mice showed the highest discrimination ratio towards the novel object (M=0.2396, p=0.0042, Fig 5F), and the pooled female group also showed a significant discrimination ratio (M=0.1155, p=0.009).

## 4. Discussion

Across all tests performed (EPM, EZM, Y-Maze, and OFT), the most robust effect observed was a reduction in exploration on repeated exposure to the apparatus. This effect was much stronger and more consistent than that of sex or circadian rhythm. This reduction in exploration is characterized by reduced locomotion and reduced time spent in threatening areas of the apparatus, such as the open arms in the EPM and the center of the open field. These findings highlight the importance of accounting for this drop in locomotion when planning studies that involve within-subject designs and provide important insight into which assays are repeatable versus which are not. The drop in locomotion interacts with the standard endpoints differently across assays. In the EPM, the decline in locomotion on the second exposure coincides with an even greater reduction in the time spent in the open arms. EZM showed a similar drop in locomotion but much less of a drop in open arm time. In SA, however, percent alternation remained stable, despite a reduction in exploration. Thus, EZM and SA are more repeatable and suitable for within-subject designs, whereas EPM is clearly not suitable (Tucker & McCabe 2017; Cnops et al. 2022; Magen et al., 2012).

We propose that the first exposure to the apparatus should not be viewed as a stable baseline, but as an incidental learning experience during which the mice form a representation of the apparatus. The initial novelty-driven exploration subsides as the environment becomes familiar and this learning is expressed on the second day as a drop in exploration. Thus, even though these assays are not intended to assess long term memory, they are inherently doing so when multiple tests are conducted in the same animal. The drop in exploration of the threatening areas of the apparatus that we observed in all three assays (EPM, EZM and OFT) suggests that the animals may also be learning about threatening areas during the initial exposure. On the second exposure, their overall reduction in exploration leads to a greater avoidance of the threatening areas. This effect is very pronounced in the EPM, such that the drop in time spent in the open arms far exceeds the overall drop in locomotion. Rather than interpreting this disproportionate avoidance of the open arms as an increase in “anxiety-like” behavior, as would be the traditional parlance, we propose that it reflects an active learning process that could be termed spatial threat learning. This theoretical reappraisal has important implications for both the design of within-subject experiments and the interpretation of the results.

In this study, we also sought to compare and investigate the joint impact of sex, circadian rhythm, and repeated testing on behavioral performance. We observed that the effects of sex and circadian phase are less robust and less consistent than the effects of a second exposure to an assay. Two tests, the EZM and Y-maze, were unaffected by either sex or reversed light cycle, indicating that these assays may be the most resilient to changing experimental variables. We did not observe any significant effects of sex in any assay, though there was a trend towards reduced closed arm activity in males the EPM. This reduction in sex differences may stem from the use of handling tunnels; while these tools have been shown to minimize handling-induced anxiety that may confound with behavioral assays, they also consequently reduce sex differences in behavior (Churchill et al., 2026).

Light cycle, on the other hand, affected the OFT and NOR assays. In the OFT, RLD mice showed increased locomotion, which indicates heightened motivation to explore. Despite this increase in exploration, RLD mice showed reduced time in the center compared to the periphery. This result could be due to improved encoding of the center as threatening, consistent with the theory that RLD improves spatial learning in the OFT (Krasne et al., 2015; Wagatsuma et al., 2018). These data warrant future research on the difference in spatial learning between RLD and SLD mice. In the NOR assay, RLD mice spent more time overall investigating the objects compared to SLD mice, especially for males on the familiarization day. This result could be explained by either increased motivation towards exploring novelty or by better detection of novelty. We observed that this increase in exploration during familiarization predicted better performance on the test day, and RLD males performed the best at identifying the novel object compared to any other group. Together, these data suggest that the improved novel object recognition performance is mediated by increased investigation of the object during the initial familiarization phase. Moreover, to improve performance and reduce variation of the NOR, thereby requiring fewer animals, RLD mice may be the best choice when running NOR.

Overall, we identify repeated testing as a robust extrinsic variable, more so than sex or circadian phase, that greatly impacts exploratory behavior in all assays, but affects typically measured endpoints in the EPM, EZM, and OFT. We hypothesize that this effect occurs due to an incidental encoding of the novel behavioral assay, which is expressed as learning upon a second exposure via a reduction in exploration. Future research should attempt to further characterize this effect, its underlying mechanisms, and how it may interact with treatment effects. For example, the 5XFAD Alzheimer’s disease mouse model shows persistently elevated exploratory behavior across a 30-minute open field (Perl et al., 2025) and fails to reduce its exploration with repeated exposure (Forner et al., 2021). Furthermore, 5XFAD mice have increased open arm time in the EPM, which is typically interpreted as decreased anxiety-like behavior (Flanigan et al., 2014). Both effects might possibly reflect the same underlying deficit in incidental learning : the environment stays persistently novel thereby driving aberrantly elevated exploratory behavior in the OFT and repeated ventures into the more threatening open arms in the EPM (Smith & Hopp, 2023). In addition, we find that EPM is affected by sex, OFT and NOR are affected by a reversed light cycle, and EZM and Y-maze are not affected by sex or reversed light cycles. In this study, we define the limits of interpretability across experiments which differ in these methodological choices. These results also serve to inform experimental design choices for future neurobehavioral studies, as our results show the behavioral effects of repeated testing are significant, and warrant further investigation of these effects as well as the learning processes that may be causing them.

## CRediT authorship contribution statement

**Jake E. Riggi:** Conceptualization, Methodology, Formal Analysis, Investigation, Writing – Original Draft, Writing - Review & Editing. **Margaret H. Copeland:** Conceptualization, Methodology, Formal Analysis, Writing - Original Draft, Writing - Review & Editing, Visualization, Supervision, Project administration. **Sydney J. Schaefer:** Conceptualization, Methodology, Validation, Formal Analysis, Investigation, Writing - Review & Editing. **Maya Yun:** Methodology, Formal Analysis, Writing - Original Draft, Writing - Review & Editing. **Jesse D. Cushman:** Conceptualization, Methodology, Writing - Review & Editing, Resources, Funding acquisition, Supervision.

## Acknowledgements

We thank Korey D. Stevanovic and Leslie R. Aksu for their expert guidance on behavioral testing techniques.

## Data availability

Data will be made available upon request.

## Declaration of generative AI and AI-assisted technologies in the manuscript preparation process

During the preparation of this work, the authors used ChatGPT-6.1 tools to assist with the literature review, edit behavioral diagrams, and edit code to format plots. The authors reviewed and edited the output as needed and take full responsibility for the content of the published article.

## References

Albernaz-Mariano, K. A., Souza, R. R., & Canto-de-Souza, A. (2022). Blockade of the mineralocorticoid receptors in the medial prefrontal cortex prevents the acquisition of one-trial tolerance in mice. Behavioural Brain Research, 431, 113938. 10.1016/j.bbr.2022.113938

Almeida, A., Tavares, I., Lima, D., & Coimbra, A. (1993). Descending projections from the medullary dorsal reticular nucleus make synaptic contacts with spinal cord lamina I cells projecting to that nucleus: an electron microscopic tracer study in the rat. Neuroscience, 55(4), 1093–1106. 10.1016/0306-4522(93)90323-8

Alstott, J., & Timberlake, W. (2009). Effects of rat sex differences and lighting on locomotor exploration of a circular open field with free-standing central corners and without peripheral walls. Behavioural Brain Research, 196(2), 214–219. 10.1016/j.bbr.2008.09.001

Baumgartner, N. E., Biraud, M. C., & Lucas, E. K. (2023). Sex differences in socioemotional behavior and changes in ventral hippocampal transcription across aging in C57Bl/6J mice. Neurobiology of Aging, 130, 141–153. 10.1016/j.neurobiolaging.2023.05.015

Beery, A. K., & Zucker, I. (2011). Sex bias in neuroscience and biomedical research. Neuroscience and Biobehavioral Reviews, 35(3), 565–572. 10.1016/j.neubiorev.2010.07.002

Bernstein, B. J., Kendricks, D. R., Fry, S., Wilson, L., Koopmans, B., Loos, M., Stevanovic, K. D., & Cushman, J. D. (2024). Sex differences in spontaneous behavior and cognition in mice using an automated behavior monitoring system. Physiology & Behavior, 283, 114595. 10.1016/j.physbeh.2024.114595

Bertoglio, L. J., & Carobrez, A. P. (2000). Previous maze experience required to increase open arms avoidance in rats submitted to the elevated plus-maze model of anxiety. Behavioural Brain Research, 108(2), 197–203. 10.1016/s0166-4328(99)00148-5

Chen, W., Wang, Z., Ma, C., Ma, X., Meng, W., Yin, F., & Yang, Y. (2023). Tactile cues are important to environmental novelty during repeated open field tests. Behavioural Processes, 204, 104796. 10.1016/j.beproc.2022.104796

Churchill, S. R., Stallone, J., Aksu, L., Webb, D. R., King-Herbert, A., Cushman, J. D., Churchill, S. R., Stallone, J., Aksu, L., Webb, D. R., King-Herbert, A., & Cushman, J. D. (2026). Refined Handling Reduces Anxiety-Related Behavior and Improves Object Recognition Performance in C57BL/6J Mice. Journal of the American Association for Laboratory Animal Science, 1.

Clayton, J. A., & Collins, F. S. (2014). Policy: NIH to balance sex in cell and animal studies. Nature, 509(7500), 282–283. 10.1038/509282a

Cnops, V., Iyer, V. R., Parathy, N., Wong, P., & Dawe, G. S. (2022). Test, rinse, repeat: A review of carryover effects in rodent behavioral assays. Neuroscience and Biobehavioral Reviews, 135, 104560. 10.1016/j.neubiorev.2022.104560

Cook, M. N., Crounse, M., & Flaherty, L. (2002). Anxiety in the elevated zero-maze is augmented in mice after repeated daily exposure. Behavior Genetics, 32(2), 113–118. 10.1023/a:1015249706579

Crawley, J. N. (2008). Behavioral phenotyping strategies for mutant mice. Neuron, 57(6), 809–818. 10.1016/j.neuron.2008.03.001

Djebari, S., Contreras, A., Castro-Andrés, V., Jimenez-Herrera, R., Iborra-Lázaro, G., Sánchez-Campusano, R., Jiménez-Díaz, L., & Navarro-López, J. D. (2026). Posterior parietal cortex oscillatory activity reflects persistent spatial memory impairments induced by early hippocampal amyloidosis in male mice. The Journal of Physiology, 604(7), 3053–3076. 10.1113/JP286196

Fanselow, M S. (2000). Contextual fear, gestalt memories, and the hippocampus. Behavioural Brain Research, 110(1–2), 73–81. 10.1016/s0166-4328(99)00186-2

Fanselow, Michael S. (1990). Factors governing one-trial contextual conditioning. Animal Learning & Behavior, 18(3), 264–270. 10.3758/BF03205285

Ferreira, J. S., Leite Junior, J. B., de Mello Bastos, J. M., Samuels, R. I., Carey, R. J., & Carrera, M. P. (2022). A new method to study learning and memory using spontaneous locomotor activity in an open-field arena. Journal of Neuroscience Methods, 366, 109429. 10.1016/j.jneumeth.2021.109429

Festing, M. F. W., & Altman, D. G. (2002). Guidelines for the design and statistical analysis of experiments using laboratory animals. ILAR Journal / National Research Council, Institute of Laboratory Animal Resources, 43(4), 244–258. 10.1093/ilar.43.4.244

File, S. E. (1993). The interplay of learning and anxiety in the elevated plus-maze. Behavioural Brain Research, 58(1–2), 199–202. 10.1016/0166-4328(93)90103-w

File, S. E., Mabbutt, P. S., & Hitchcott, P. K. (1990). Characterisation of the phenomenon of “one-trial tolerance” to the anxiolytic effect of chlordiazepoxide in the elevated plus-maze. Psychopharmacology, 102(1), 98–101. 10.1007/BF02245751

Flanigan, T. J., Xue, Y., Kishan Rao, S., Dhanushkodi, A., & McDonald, M. P. (2014). Abnormal vibrissa-related behavior and loss of barrel field inhibitory neurons in 5xFAD transgenics. Genes, Brain, and Behavior, 13(5), 488–500. 10.1111/gbb.12133

Forner, S., Kawauchi, S., Balderrama-Gutierrez, G., Kramár, E. A., Matheos, D. P., Phan, J., Javonillo, D. I., Tran, K. M., Hingco, E., da Cunha, C., Rezaie, N., Alcantara, J. A., Baglietto-Vargas, D., Jansen, C., Neumann, J., Wood, M. A., MacGregor, G. R., Mortazavi, A., Tenner, A. J., … Green, K. N. (2021). Systematic phenotyping and characterization of the 5xFAD mouse model of Alzheimer’s disease. Scientific Data, 8(1), 270. 10.1038/s41597-021-01054-y

Hånell, A., & Marklund, N. (2014). Structured evaluation of rodent behavioral tests used in drug discovery research. Frontiers in Behavioral Neuroscience, 8, 252. 10.3389/fnbeh.2014.00252

Heald, J. B., Wolpert, D. M., & Lengyel, M. (2023). The Computational and Neural Bases of Context-Dependent Learning. Annual Review of Neuroscience, 46, 233–258. 10.1146/annurev-neuro-092322-100402

Hughes, R. N. (2004). The value of spontaneous alternation behavior (SAB) as a test of retention in pharmacological investigations of memory. Neuroscience and Biobehavioral Reviews, 28(5), 497–505. 10.1016/j.neubiorev.2004.06.006

Karp, N. A., Mason, J., Beaudet, A. L., Benjamini, Y., Bower, L., Braun, R. E., Brown, S. D. M., Chesler, E. J., Dickinson, M. E., Flenniken, A. M., Fuchs, H., Angelis, M. H. de, Gao, X., Guo, S., Greenaway, S., Heller, R., Herault, Y., Justice, M. J., Kurbatova, N., … White, J. K. (2017). Prevalence of sexual dimorphism in mammalian phenotypic traits. Nature Communications, 8, 15475. 10.1038/ncomms15475

Kitamura, T., Okubo-Suzuki, R., Takashima, N., Murayama, A., Hino, T., Nishizono, H., Kida, S., & Inokuchi, K. (2012). Hippocampal function is not required for the precision of remote place memory. Molecular Brain, 5, 5. 10.1186/1756-6606-5-5

Kraeuter, A.-K., Guest, P. C., & Sarnyai, Z. (2019). The Y-Maze for Assessment of Spatial Working and Reference Memory in Mice. Methods in Molecular Biology, 1916, 105–111. 10.1007/978-1-4939-8994-2_10

Krasne, F. B., Cushman, J. D., & Fanselow, M. S. (2015). A Bayesian context fear learning algorithm/automaton. Frontiers in Behavioral Neuroscience, 9, 112. 10.3389/fnbeh.2015.00112

Lalonde, R. (2002). The neurobiological basis of spontaneous alternation. Neuroscience and Biobehavioral Reviews, 26(1), 91–104. 10.1016/s0149-7634(01)00041-0

Leussis, M. P., & Bolivar, V. J. (2006). Habituation in rodents: a review of behavior, neurobiology, and genetics. Neuroscience and Biobehavioral Reviews, 30(7), 1045–1064. 10.1016/j.neubiorev.2006.03.006

Lister, R. G. (1987). The use of a plus-maze to measure anxiety in the mouse. Psychopharmacology, 92(2), 180–185. 10.1007/BF00177912

Magen, I., Fleming, S. M., Zhu, C., Garcia, E. C., Cardiff, K. M., Dinh, D., De La Rosa, K., Sanchez, M., Torres, E. R., Masliah, E., Jentsch, J. D., & Chesselet, M.-F. (2012). Cognitive deficits in a mouse model of pre-manifest Parkinson’s disease. The European Journal of Neuroscience, 35(6), 870–882. 10.1111/j.1460-9568.2012.08012.x

Mariscal, P., Bravo, L., Llorca-Torralba, M., Razquin, J., Miguelez, C., Suárez-Pereira, I., & Berrocoso, E. (2023). Sexual differences in locus coeruleus neurons and related behavior in C57BL/6J mice. Biology of Sex Differences, 14(1), 64. 10.1186/s13293-023-00550-7

McIlwain, K. L., Merriweather, M. Y., Yuva-Paylor, L. A., & Paylor, R. (2001). The use of behavioral test batteries: effects of training history. Physiology & Behavior, 73(5), 705–717. 10.1016/s0031-9384(01)00528-5

Nakano, J. J., Shimizu, K., Shimba, S., & Fukada, Y. (2016). SCOP/PHLPP1β in the basolateral amygdala regulates circadian expression of mouse anxiety-like behavior. Scientific Reports, 6, 33500. 10.1038/srep33500

Nedogreeva, O. A., Недогреева О., А., Manolova, A. O., Манолова А., О., Mamedova, D. I., Мамедова Д., И., Gulyaeva, N. V., Гуляева Н., В., Stepanichev, M. Y., & Степаничев М., Ю. (2025). Non-associative learning (habituation) in normotensive and hypertensive rats and the effects of social isolation. I.P. Pavlov Journal of Higher Nervous Activity, 75.

O’Reilly, R. C., & Rudy, J. W. (2001). Conjunctive representations in learning and memory: principles of cortical and hippocampal function. Psychological Review, 108(2), 311–345. 10.1037/0033-295x.108.2.311

Peirson, S. N., Brown, L. A., Pothecary, C. A., Benson, L. A., & Fisk, A. S. (2018). Light and the laboratory mouse. Journal of Neuroscience Methods, 300, 26–36. 10.1016/j.jneumeth.2017.04.007

Perl, A. T., Wu, J., Dong, J. D., Brooks, A. M., Yoblinski, A. R., Vierling, T. T., Li, J.-L., Ruby, D. R., Radzicki, D., Dudek, S. M., Cushman, J. D., & Gjoneska, E. (2026). Immune receptor LAG3 regulates microglia function during Alzheimer’s disease. Neurobiology of Disease, 226, 107447. 10.1016/j.nbd.2026.107447

Pignatelli, M., Ryan, T. J., Roy, D. S., Lovett, C., Smith, L. M., Muralidhar, S., & Tonegawa, S. (2019). Engram cell excitability state determines the efficacy of memory retrieval. Neuron, 101(2), 274–284.e5. 10.1016/j.neuron.2018.11.029

Richetto, J., Polesel, M., & Weber-Stadlbauer, U. (2019). Effects of light and dark phase testing on the investigation of behavioural paradigms in mice: Relevance for behavioural neuroscience. Pharmacology, Biochemistry, and Behavior, 178, 19–29. 10.1016/j.pbb.2018.05.011

Ripperger, J. A., Jud, C., & Albrecht, U. (2011). The daily rhythm of mice. FEBS Letters, 585(10), 1384– 1392. 10.1016/j.febslet.2011.02.027

Roy, V., Chapillon, P., Jeljeli, M., Caston, J., & Belzung, C. (2009). Free versus forced exposure to an elevated plus-maze: evidence for new behavioral interpretations during test and retest. Psychopharmacology, 203(1), 131–141. 10.1007/s00213-008-1378-2

Ryherd, G. L., Bunce, A. L., Edwards, H. A., Baumgartner, N. E., & Lucas, E. K. (2023). Sex differences in avoidance behavior and cued threat memory dynamics in mice: Interactions between estrous cycle and genetic background. Hormones and Behavior, 156, 105439. 10.1016/j.yhbeh.2023.105439

Schrader, A. J., Taylor, R. M., Lowery-Gionta, E. G., & Moore, N. L. T. (2018). Repeated elevated plus maze trials as a measure for tracking within-subjects behavioral performance in rats (Rattus norvegicus). Plos One, 13(11), e0207804. 10.1371/journal.pone.0207804

Schröder, D., Wrona, K. J., Müller, F., Heinemann, S., Fischer, F., & Dockweiler, C. (2023). Impact of virtual reality applications in the treatment of anxiety disorders: A systematic review and meta-analysis of randomized-controlled trials. Journal of Behavior Therapy and Experimental Psychiatry, 81, 101893. 10.1016/j.jbtep.2023.101893

Seibenhener, M. L., & Wooten, M. C. (2015). Use of the Open Field Maze to measure locomotor and anxiety-like behavior in mice. Journal of Visualized Experiments, 96, e52434. 10.3791/52434

Sergievich, A. A., Khoroshikh, P. P., Artemenko, A. F., Zakharenko, A. M., Chaika, V. V., Kodintsev, V. V., Stroeva, O. A., Lenda, E. G., Tsatsakis, A., Burykina, T. I., Agathokleous, E., Kostoff, R. N., Zlatian, O., Docea, A. O., & Golokhvast, K. S. (2020). Behavioral impacts of a mixture of six pesticides on rats. The Science of the Total Environment, 727, 138491. 10.1016/j.scitotenv.2020.138491

Smith, A. J., & Lilley, E. (2019). The role of the three rs in improving the planning and reproducibility of animal experiments. Animals : An Open Access Journal from MDPI, 9(11). 10.3390/ani9110975

Smith, S., & Hopp, S. C. (2023). The 5XFAD mouse model of Alzheimer’s disease displays age-dependent deficits in habituation to a novel environment. Aging Brain, 3, 100078. 10.1016/j.nbas.2023.100078

Sousa, N., Almeida, O. F. X., & Wotjak, C. T. (2006). A hitchhiker’s guide to behavioral analysis in laboratory rodents. Genes, Brain, and Behavior, 5 Suppl 2, 5–24. 10.1111/j.1601-183X.2006.00228.x

Stevanovic, K. D., Fry, S. A., DeFilipp, J. M. S., Wu, N., Bernstein, B. J., & Cushman, J. D. (2022). Assessing the importance of sex in a hippocampus-dependent behavioral test battery in C57BL/6NTac mice. Learning & Memory, 29(8), 203–215. 10.1101/lm.053599.122

Tolman, E. C. (1948). Cognitive maps in rats and men. Psychological Review, 55(4), 189–208. 10.1037/h0061626

Tsao, C.-H., Flint, J., & Huang, G.-J. (2022). Influence of diurnal phase on behavioral tests of sensorimotor performance, anxiety, learning and memory in mice. Scientific Reports, 12(1), 432. 10.1038/s41598-021-03155-5

Tucker, L. B., & McCabe, J. T. (2017). Behavior of Male and Female C57BL/6J Mice Is More Consistent with Repeated Trials in the Elevated Zero Maze than in the Elevated Plus Maze. Frontiers in Behavioral Neuroscience, 11, 13. 10.3389/fnbeh.2017.00013

Tzakis, N., Ethier-Gagnon, M., Epp, T., & Holahan, M. R. (2025). Assessment of cFos labeling in the hippocampus and anterior cingulate cortex following recent and remote re-exposure to an unreinforced open field in preadolescent and postadolescent rats. Behavioural Brain Research, 476, 115284. 10.1016/j.bbr.2024.115284

Varga, Z. K., Pejtsik, D., Tóth, M., Balogh, Z., Aliczki, M., Szente, L., Balla, G. Y., Kontra, L., Eckert, Z., Szebik, H., Borhegyi, Z., & Mikics, É. (2025). Repeated behavioral testing and the use of summary measures reveal trait anxiety in preclinical rodent models. Translational Psychiatry, 15(1), 440. 10.1038/s41398-025-03586-y

Võikar, V., Kõks, S., Vasar, E., & Rauvala, H. (2001). Strain and gender differences in the behavior of mouse lines commonly used in transgenic studies. Physiology & Behavior, 72(1–2), 271–281. 10.1016/s0031-9384(00)00405-4

Võikar, V., Vasar, E., & Rauvala, H. (2004). Behavioral alterations induced by repeated testing in C57BL/6J and 129S2/Sv mice: implications for phenotyping screens. Genes, Brain, and Behavior, 3(1), 27–38. 10.1046/j.1601-183x.2003.0044.x

Wagatsuma, A., Okuyama, T., Sun, C., Smith, L. M., Abe, K., & Tonegawa, S. (2018). Locus coeruleus input to hippocampal CA3 drives single-trial learning of a novel context. Proceedings of the National Academy of Sciences of the United States of America, 115(2), E310–E316. 10.1073/pnas.1714082115

Walf, A. A., & Frye, C. A. (2007). The use of the elevated plus maze as an assay of anxiety-related behavior in rodents. Nature Protocols, 2(2), 322–328. 10.1038/nprot.2007.44

Weber, H., Statz, M., Markert, F., Storch, A., & Fauser, M. (2024). Circadian variations influence anxiety-related behaviour, olfaction, and hedonic response in male Sprague-Dawley rats. Behavioural Brain Research, 471, 115134. 10.1016/j.bbr.2024.115134

Whittington, J. C. R., Muller, T. H., Mark, S., Chen, G., Barry, C., Burgess, N., & Behrens, T. E. J. (2020). The Tolman-Eichenbaum Machine: Unifying Space and Relational Memory through Generalization in the Hippocampal Formation. Cell, 183(5), 1249–1263.e23. 10.1016/j.cell.2020.10.024

Young, P. A., Waller, O., Ball, K., Williams, C. C., & Nashmi, R. (2024). Phasic stimulation of dopaminergic neurons of the lateral substantia nigra increases open field exploratory behaviour and reduces habituation over time. Neuroscience, 551, 276–289. 10.1016/j.neuroscience.2024.05.025

