## Supplementary figures and images for "Prior Experience Alters Exploration in Standard Behavioral Assays More Robustly Than Sex or Light/Dark Cycle in C57BL/6J Mice"

### Supplement 1

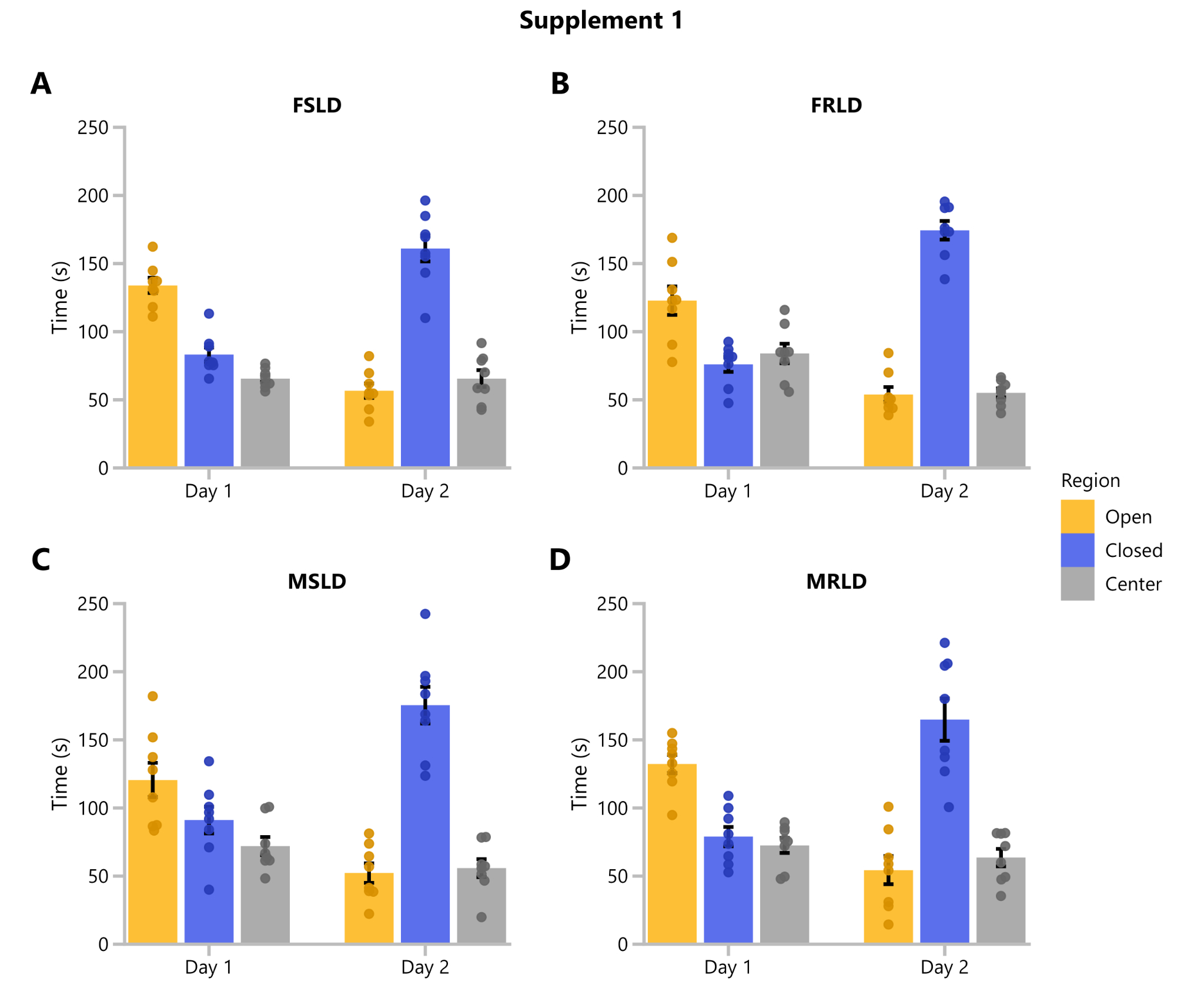

### Supplement 2

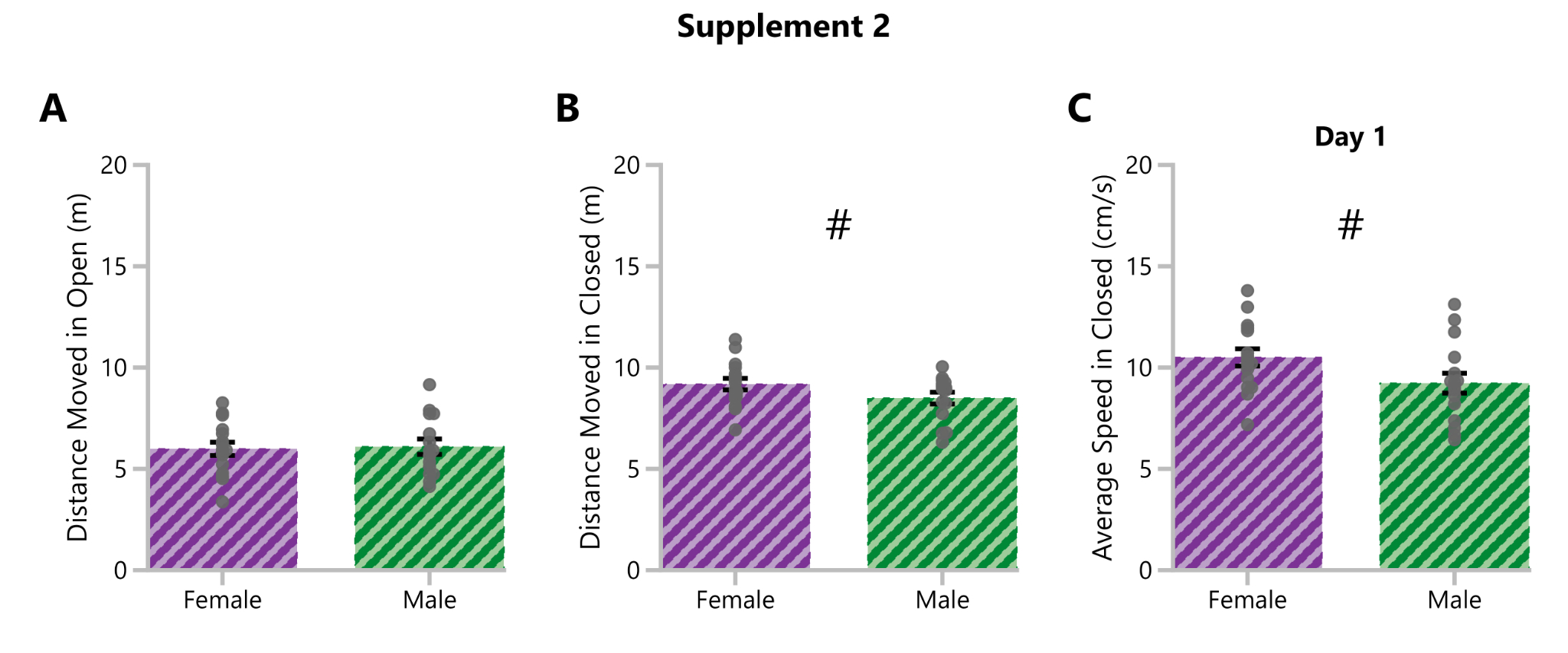

### Supplement 3

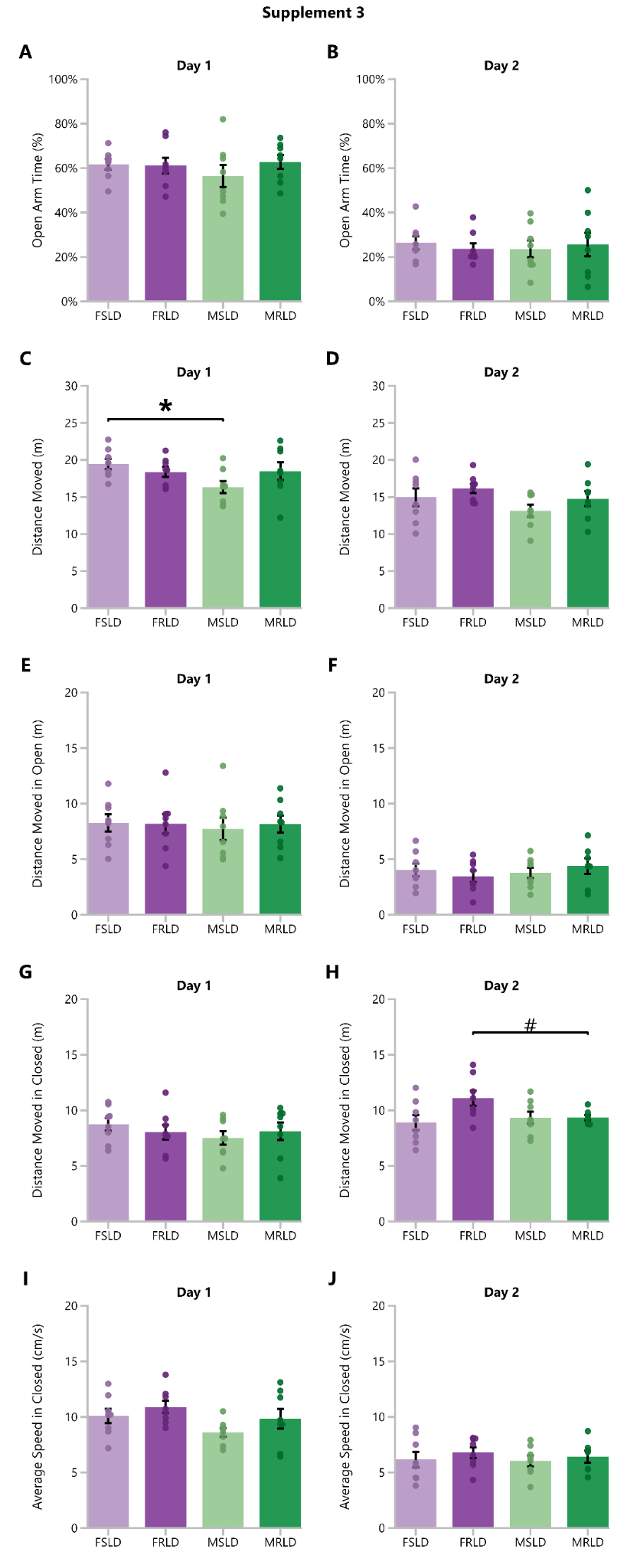

### Supplement 4

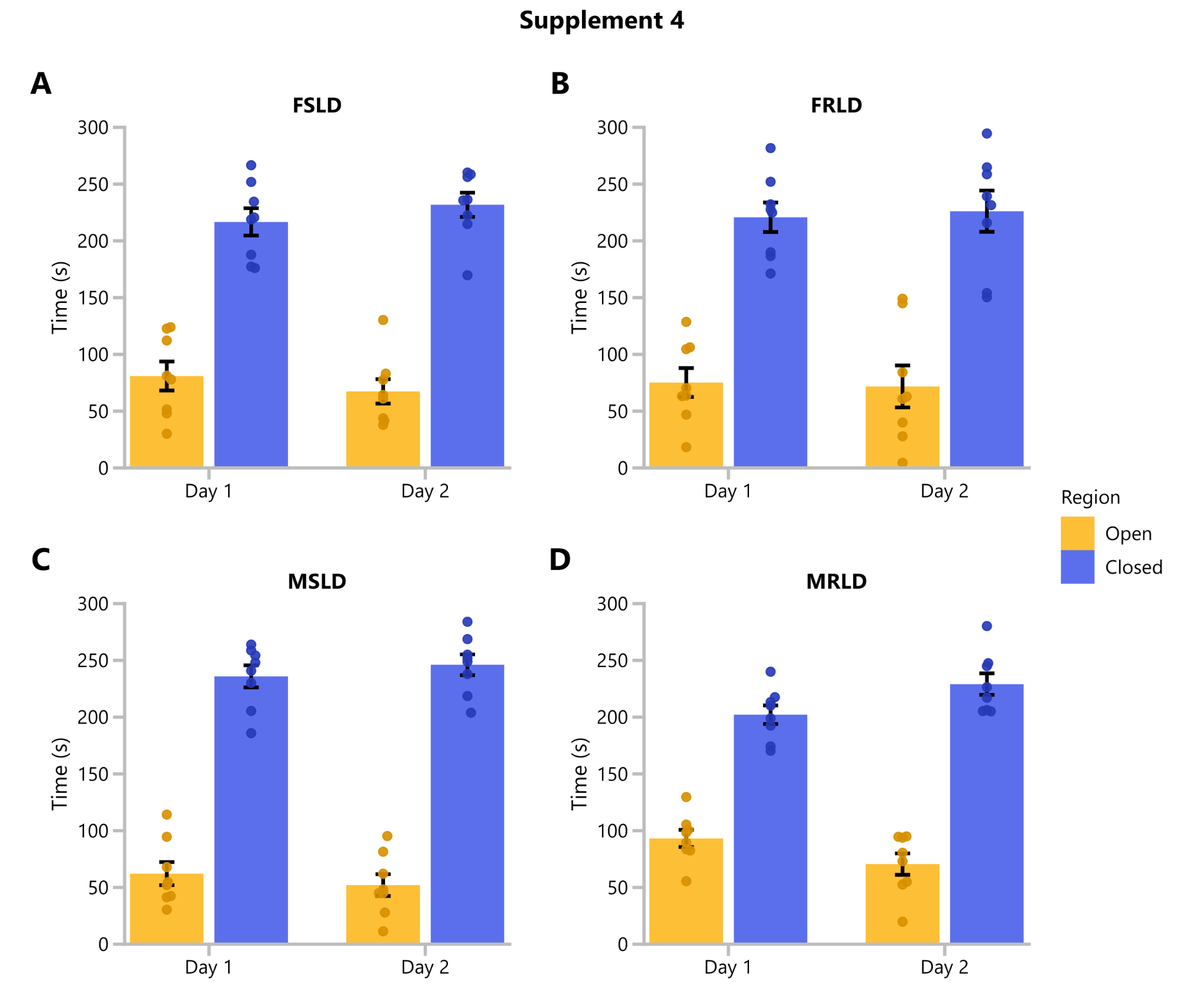

### Supplement 5

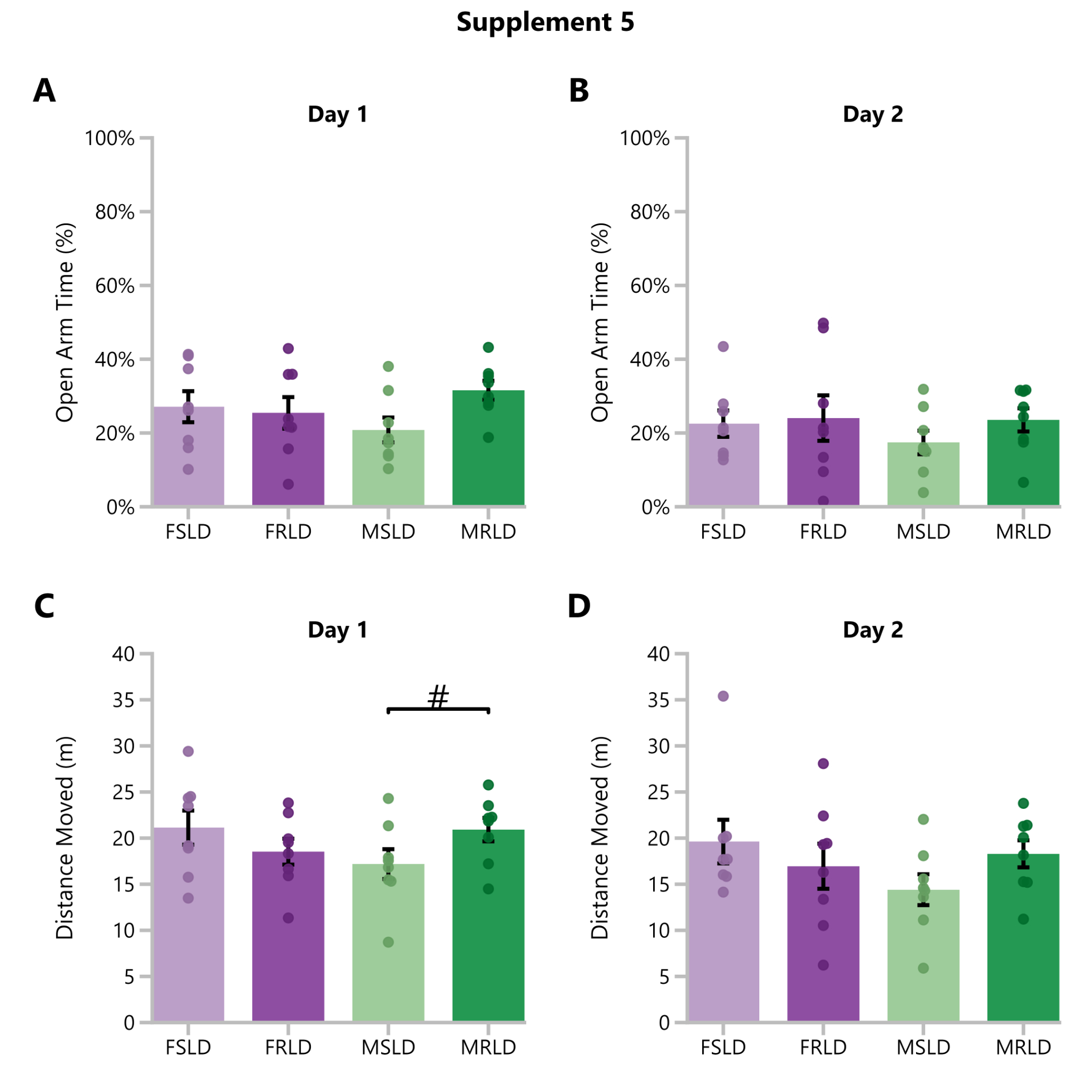

### Supplement 6

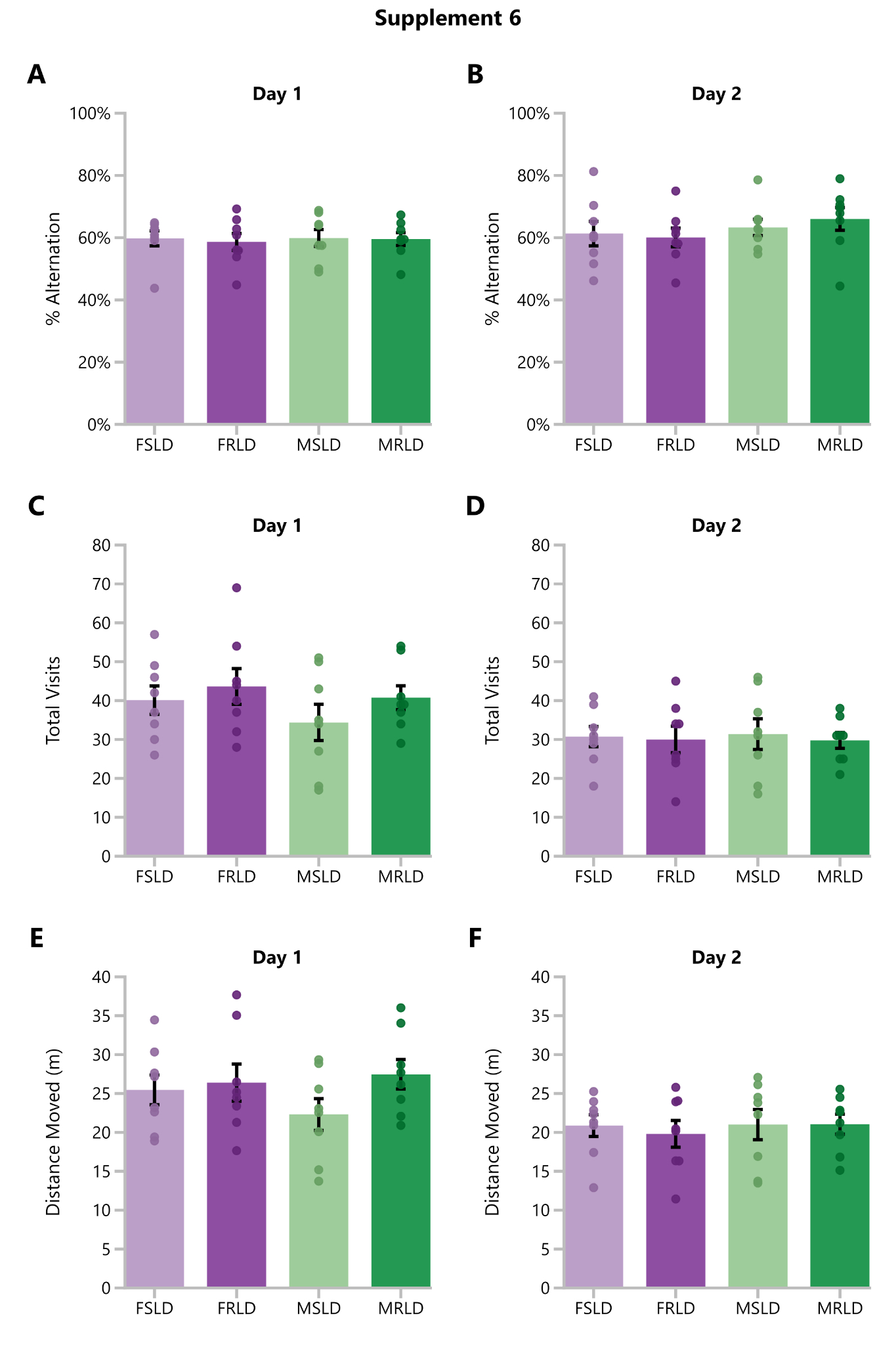

### Supplement 7

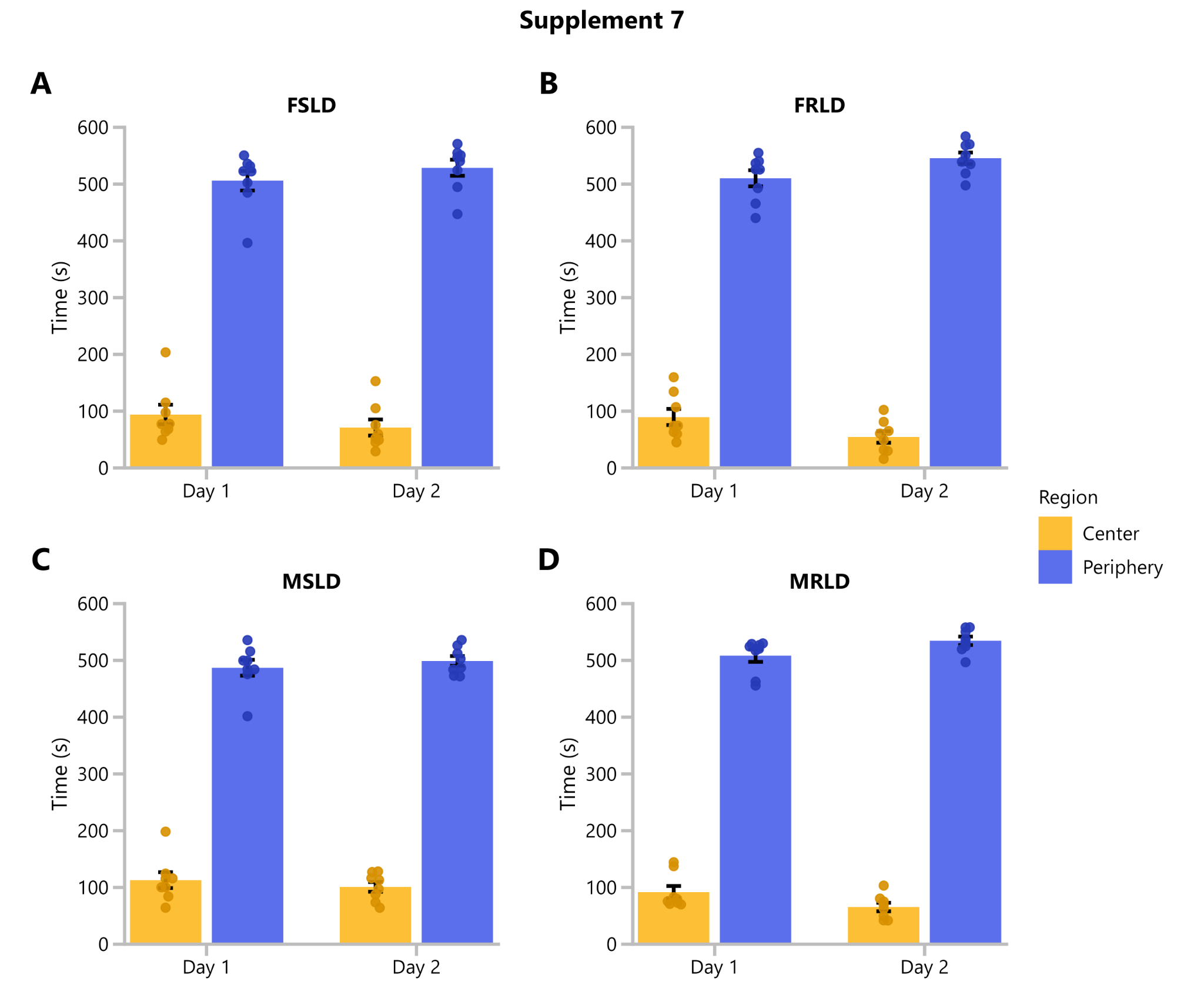

### Supplement 8

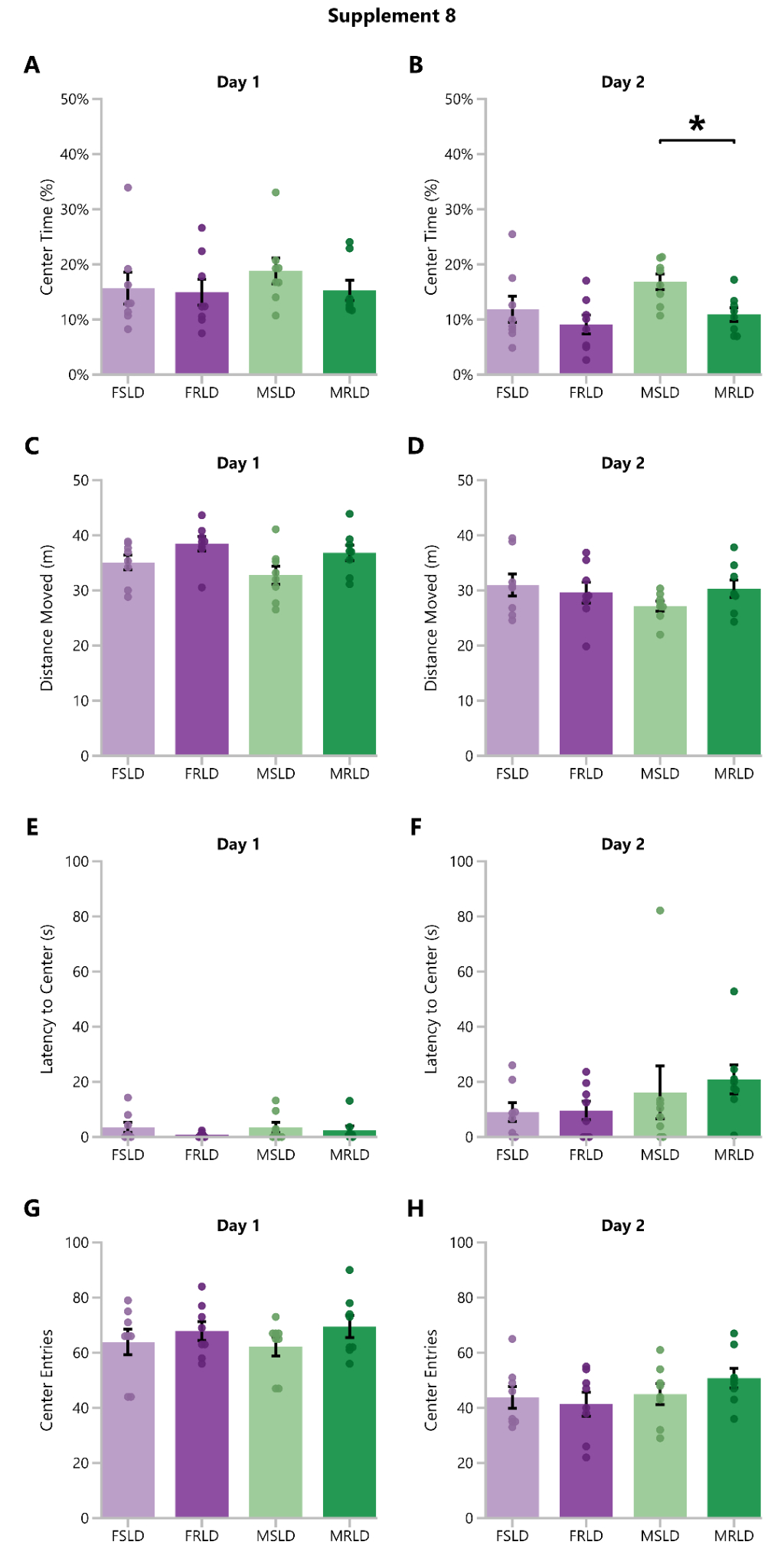

### Supplement 9

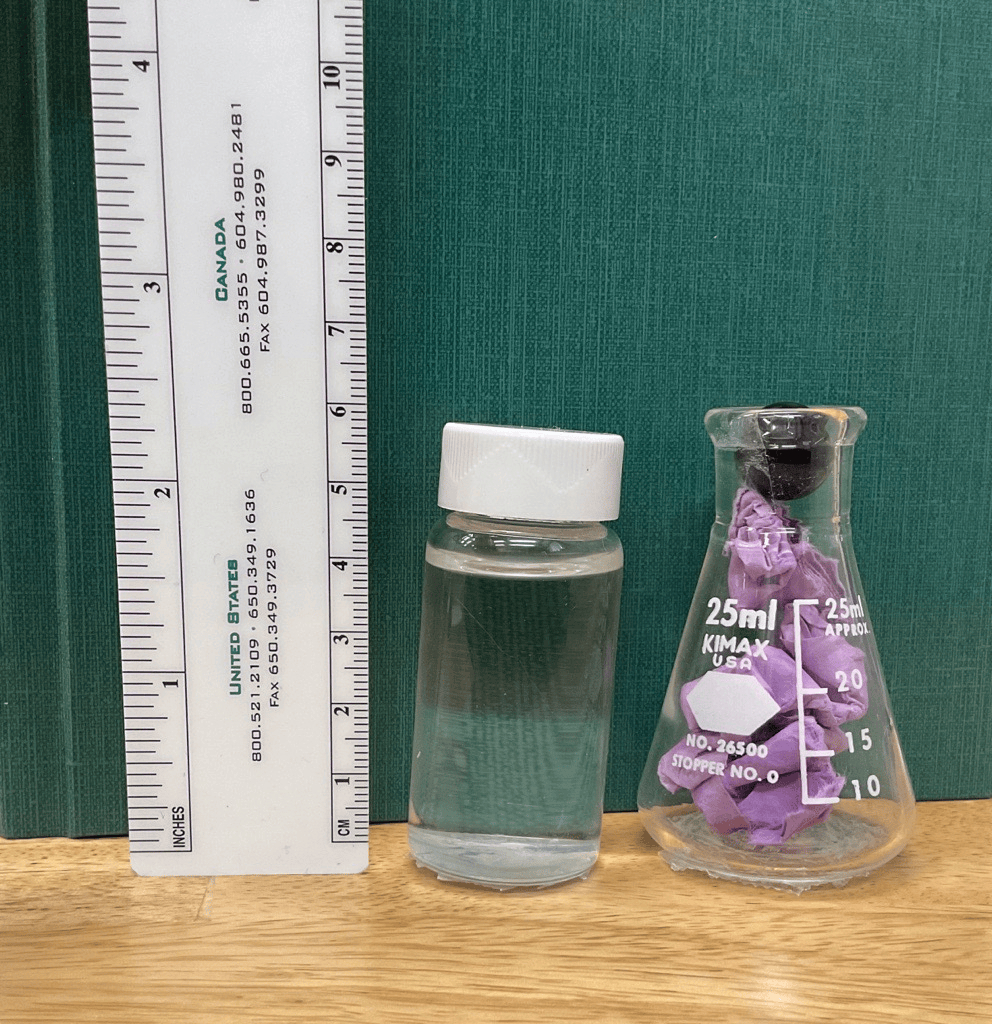

### Supplement 10

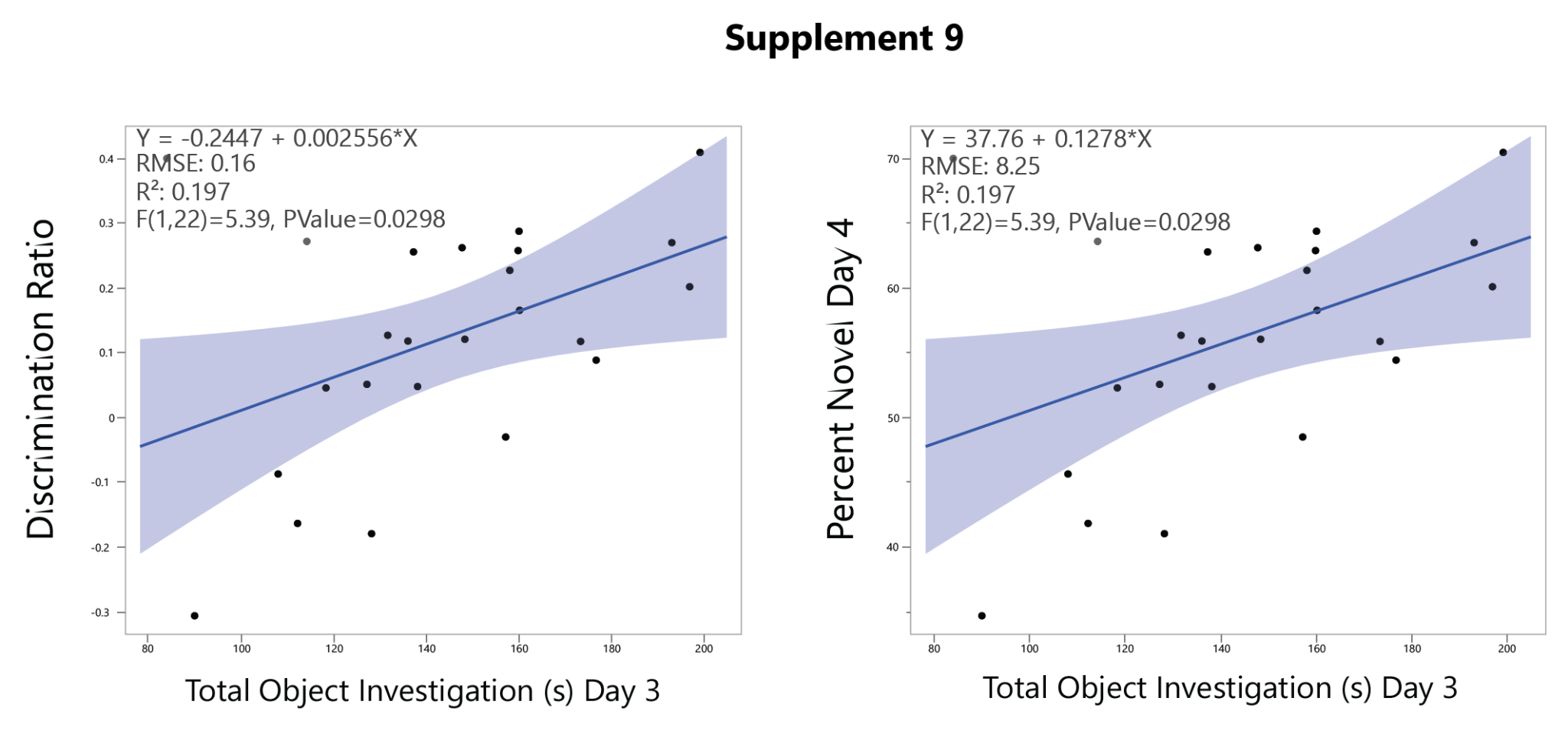
